# Deducing splicing events from transcript sets with splicelogic

**DOI:** 10.64898/2026.10.30.762958

**Authors:** Beatriz Campillo, Michael I. Love

**Affiliations:** Bioinformatics and Computational Biology Curriculum, University of North Carolina at Chapel Hill; Department of Genetics, University of North Carolina at Chapel Hill; Department of Biostatistics, University of North Carolina at Chapel Hill

**Keywords:** alternative splicing, differential isoform usage, differential transcript usage, long-read RNA-seq

## Abstract

Long-read RNA sequencing captures a greater proportion of full-length transcripts, enabling more accurate isoform identification, discovery of novel isoforms, and more precise differential transcript usage analysis. However, comparing transcripts, whether between conditions or between sets such as novel and annotated isoforms, yields complex isoform changes that are hard to manipulate, summarize, and visualize. This is compounded by vertical integration of some tools, where downstream analysis depends on the specific output of a particular upstream tool. To address this gap, we have developed splicelogic, an R/Bioconductor package that makes transcript-set comparisons biologically interpretable by turning them into discrete splicing events, such as exon skipping, alternative splice sites, intron retention, and alternative transcription start and end sites. We designed splicelogic to integrate easily with existing RNA-seq workflows and tools. The splicelogic functions are modular, readable, and extensible, making it easy for users to adapt core logic to their own custom scientific questions or novel splicing definitions. By obtaining interpretable splicing events from transcript changes, splicelogic enables downstream analyses, including annotation of RNA-binding protein recognition sites, alongside summary statistics and visualization. The splicelogic package is publicly available and distributed through Bioconductor.

## Introduction

A single gene can give rise to multiple distinct mRNA isoforms through alternative transcription start sites, alternative splicing, and alternative transcription end sites. Splicing removes introns from the pre-mRNA and joins exons to form the mature transcript. Alternative promoters and polyadenylation sites change the 5′ and 3′ ends of the transcript, altering UTRs and, in some cases, the first or last coding exons. Together these processes are a major mechanism of gene regulation and a key source of protein diversity, and isoform usage can shift in response to different conditions or over time.

Despite the central role of splicing in transcriptome complexity, the systematic detection and functional interpretation of splicing changes remain a significant challenge. Interpreting these changes requires understanding both what drives them, such as changes in RNA-binding protein activity, and what they lead to, such as altered protein function, RNA localization, or transcript stability, including nonsense-mediated decay. Splicing changes between conditions, tissues, or cell types are rarely a single event: one condition may favor skipping of some exons and inclusion of others, while also shifting splice sites, retaining introns, or switching first and last exons. Summarizing these directional shifts across many genes and event types remains difficult.

RNA-seq has become the standard tool for quantifying iso-form expression. Short-read RNA-seq is cost-effective and widely available. However, because most short reads are compatible with multiple overlapping transcripts, assigning them to specific isoforms is inherently ambiguous. Short-read sequencing limits resolution of complex structures involving multiple splicing events or alternative UTRs, as well as novel transcripts. As a result, most splicing detection methods for short reads work at the level of individual events, quantifying splicing as ratios of junction-spanning read counts without the context of the full transcript.

Beyond junction-level methods, several tools work with whole transcripts. *SUPPA2* [1] and *IsoformSwitchAnalyzeR* [2] accept abundances from common quantifiers but apply their own differential tests, so neither tool can take results from an arbitrary differential transcript usage (DTU) method. *GeneStructureTools* [3] interprets events only from junction-based tools, and long-read tools such as *SQANTI3* [4] and *IsoQuant* [5] classify novel isoforms relative to the reference rather than calling directional events between transcript sets. *FLAIR* [6] derives intron retention, alternative 5′ and 3′ splice site, and exon skipping events from isoform models assembled by *FLAIR* and read counts from its own quantification step, and tests their differential usage. It also predicts isoform productivity from open reading frames. *Swan* [7] is flexible, accepting any transcriptome as a GTF together with a transcript abundance table. It tracks the usage of transcripts and of transcription start and end sites, and, when paired with a companion tool, of intron chains. *Swan* tests each of these for switching between two conditions with its own differential test, and can use a splice graph to detect exon skipping, intron retention, and alternative splice site events relative to the reference annotation.

Long-read RNA sequencing captures a greater proportion of full-length transcripts, enabling more accurate isoform identification, discovery of novel isoforms, and more precise DTU analysis. However, the isoform changes underlying shifts in usage are complex, and asking questions outside a tool’s predefined analyses typically requires custom code that may be difficult for others to read and adapt. This is compounded by the vertical integration of some existing tools, where each downstream analysis depends on specific output of a particular upstream tool.

Here we present `splicelogic,` an R/Bioconductor [8] package that turns any two sets of transcripts into discrete splicing events, working equally well with results from any DTU method or with any user-defined partition of the transcripts. We designed `splicelogic` as a modular, efficient, and flexible framework that integrates easily with existing RNA-seq workflows and tools, and allows customized downstream computation and visualization.

### Approach

#### 1. Overview

Given two contrasting sets of transcripts, we describe their differences as a series of discrete splicing events. Transcript sets come either from a user-supplied partition or from a DTU analysis, in which each transcript with a detected change has a signed effect estimate such as a model coefficient, a change in isoform fraction, or Δ*ψ* (change in percent spliced in). Because the input is a set of transcript-level effect estimates, any upstream DTU method can be used, including *DRIMSeq* [9], *DEXSeq* [10], *satuRn* [11], and *edgeR* [12].

Event detection is built on vectorized integer-range operations from *GenomicRanges* [13] and typically takes a fraction of a second on large transcript sets from long-read RNA-seq experiments. The framework is modular and fits into existing workflows: results are projected from transcript space to the space of exon events, so users can operate directly on the detected events and their coordinates. Splice events and their consequences are represented as standard range objects, and annotations are provided as metadata columns available for computation. This allows users to directly ask subsequent questions with tools such as *plyranges* or *Biostrings*, for example to compare nucleotide frequencies around splice events or the proportions of event types.

#### 2. Input: two entry points

`splicelogic` takes as input the exon ranges for a set of transcripts across all genes, with each exon labeled by its gene, its transcript, its rank within the transcript, and a value giving the direction of change of that transcript. This input can be built from a DTU analysis or from a defined partition of the transcripts into two sets. With DTU results, the transcript annotation is combined with the per-transcript effect estimates, and the sign of each estimate marks a transcript as “positive” (up-regulated) or “negative” (down-regulated). With a set partition, such as novel versus annotated iso-forms or disease versus control, no estimates are needed: the reference set is “negative” and the alternative set is “positive”. Either route yields the same input, and from that point on the workflow is identical.

#### 3. Event detection

`splicelogic` detects various types of events: skipped exons (SE), included exons (IE), mutually exclusive exons (MXE), retained introns (RI), alternative 5′ and 3′ splice sites (A5SS, A3SS), and alternative transcription start and end sites (ATSS, ATES). Each event is defined between a pair of transcripts from the same gene, one from each group, by comparing their exon structures. For SE, IE, and MXE, neighboring exons can be matched across transcript pairs using different levels of stringency (Supplementary Methods). Each event type has its own function, which can be called by a short name or by a longer alias for readability, so `find_se()` and `find_skipped_exons()` are equivalent, and `find_all_events()` runs all of them at once.

#### 4. Output

Each `find_*()` function returns the exons involved in an event, one row per exon and paired transcript. Each row keeps the user’s original exon-level metadata, which describes the transcript that contains the returned exon, and adds event-specific columns, prefixed with `event_,` that describe the paired transcript against which the event is defined: the event type, the paired transcript’s ID and estimate, and any additional columns requested by the user (*e*.*g. ψ* per group).

#### 5. Design and integrati

Detected events are returned as genomic ranges with metadata, so downstream analysis can use standard Bioconductor tools rather than package-specific functions. For example, the nucleotide sequence of each event, or of its flanking intronic region, can be extracted with the helper function `splicelogic::get_seq()` and subsequently analyzed with *Biostrings* summarization or pattern-matching functions. Events can be manipulated or summarized with readable syntax from *plyranges* [14], part of the tidyomics ecosystem [15].

In `splicelogic` and in other splicing annotation workflows, an exon can participate in an event against more than one paired transcript, and so may appear on several rows.

Grouping by exon and event type and collapsing with *plyranges* results in one call per exon and event type, with the paired transcript IDs concatenated in metadata columns (Supplementary Code 1). This facilitates transcriptome-wide summaries of numbers of events by type, without numbers being inflated across transcript pairs within a gene. Similarly, grouping events by transcript pair identifies pairs carrying more than one event, revealing complex splicing patterns. These downstream use cases are demonstrated in the `splicelogic` vignette. Each event type is defined in a short function built from standard range operations, so users can inspect the core logic and adapt it to their own questions or to new event definitions.

## Results

### 1. Speed benchmark on simulated data

We benchmarked the speed of splicelogic on simulated data. We varied the number of genes, the number of “positive” transcripts per gene, each compared against one “negative” transcript, and the fraction of genes carrying a skipped-exon event. Runtime of `find_se()` increased linearly with the total number of exons and remained under 0.3 s for the largest case of 150,000 exons (5,000 genes, 2 transcripts per gene, and 15 exons per transcript; Figure 1a; Supplementary Methods). The other type settings of `find_se()` and event detection functions performed similarly.

**Figure 1:**
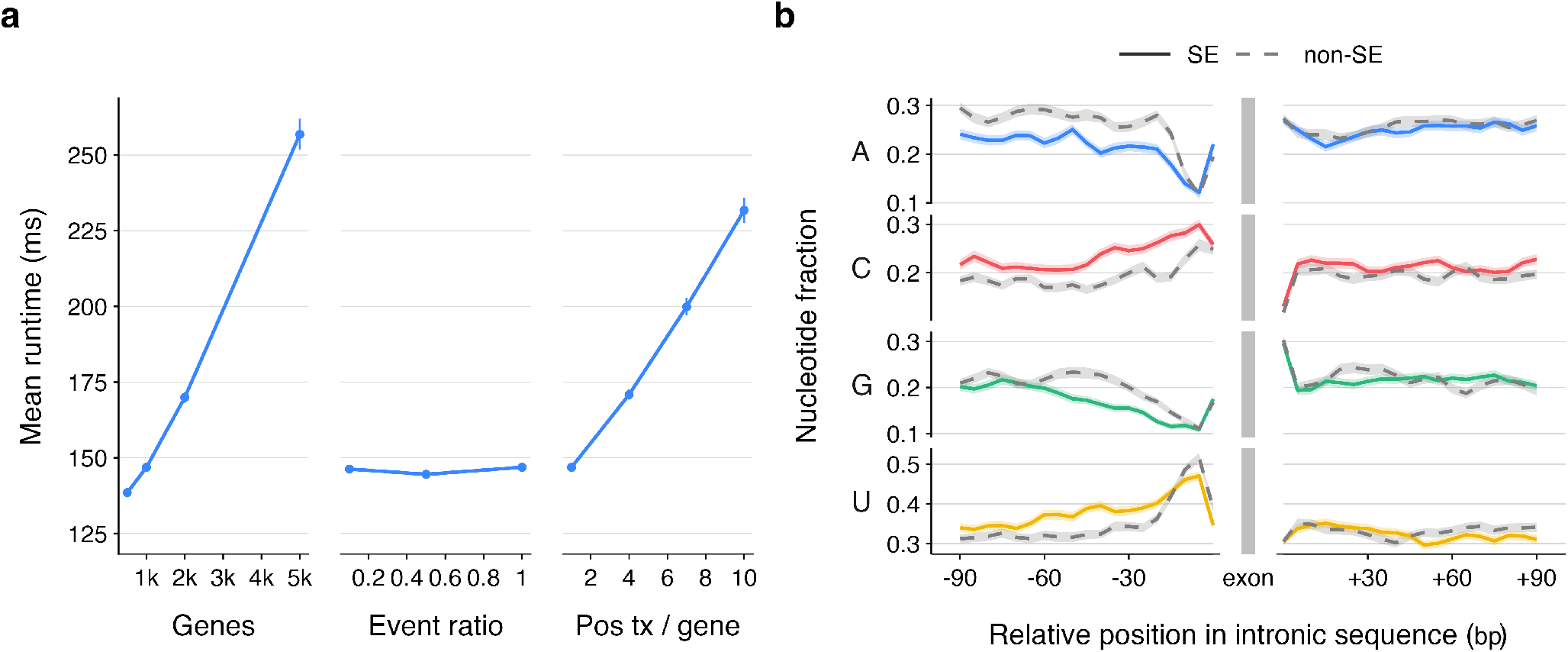
Speed of event detection and intronic context around skipped exons. (a) Runtime of `find_se()` on simulated data, shown as mean ±s.e. over 200 iterations. (b) Nucleotide composition of the intronic sequence flanking skipped exons (SE) detected by `find_se()` in transcripts with significant DTU after PTBP1 knockdown, shown as solid lines, compared to random non-skipped exons (non-SE) from transcripts without DTU, shown as dashed lines.

### 2. Correctness

We validated `splicelogic` splicing event detection against *Biosurfer* [16]. *Biosurfer* compares pairs of GENCODE protein isoforms and classifies the splicing events behind each region where their protein sequences differ. From its GEN-CODE v42 output, we selected isoform pairs that differ by a single skipped exon, included exon, alternative 5′ splice site, or alternative 3′ splice site, sampling up to 2,000 pairs per event type. We then ran the matching `splicelogic` function on each pair (Supplementary Methods). For each detected event, we translated the skipped or included exon, or the region between the two alternative splice sites, with `translate()` from *Biostrings*. For every event type, the translated amino acid sequence was identical to the sequence *Biosurfer* reported as gained or lost, both for exons in phase and for exons out of phase. We additionally sampled up to 2,000 pairs of transcripts with retained introns, where `splicelogic` detected all retained introns at the level of coordinates (Supplementary Methods).

### 3. Application datasets

The `splicelogic` vignette demonstrates the package on a published long-read dataset [17], in which Jones *et al*. used Oxford Nanopore Technologies (ONT) sequencing to compare splicing across four mouse brain regions and between sexes. We used the authors’ DTU results, focusing on female versus male cortex and splitting the 26 transcripts with FDR < 0.1 into up- and down-regulated sets by the DTU coefficient. From these, `splicelogic` recovered 22 events in 10 genes and 20 transcripts.

We also applied `splicelogic` to a long-read ONT dataset published by Glinos *et al*. [18] as part of GTEx v9. We focused on the long-read data of RNA from human skin fibroblasts subject to PTBP1 knockdown (KD) as well as controls. We quantified transcript abundance with *oarfish* [19] on the AnVIL platform [20], then performed DTU analysis using *satuRn* [11] and identified 1,987 transcripts in 1,023 genes with significant changes in usage (FDR < 0.1). `splicelogic` was used to detect splicing events in the presence of PTBP1 (WT) compared to reduced PTBP1 (KD).

Within the Bioconductor and tidyomics frameworks, we extracted the intronic sequence flanking detected skipped exons. The sequence upstream of PTBP1-induced skipped exons recapitulated PTBP1′s known binding preference (Figure 1b; Supplementary Methods). Example `splicelogic` code follows:

**Code 1:**
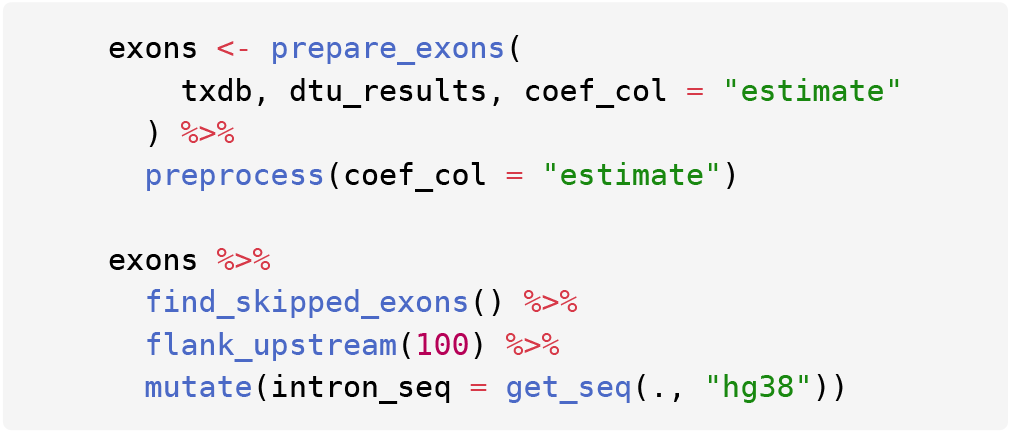
Finding skipped exons and obtaining their upstream intronic sequence. The `splicelogic` functions `find_skipped_exons()` and `get_seq()` are demonstrated within a tidyomics pipeline, including *plyranges* for range manipulation.

Lastly, we ran `splicelogic` on a transcript set comparison without DTU results, using the Glinos *et al*. dataset to compare the annotated transcripts from the GENCODE v48 annotation with the novel isoforms detected by *bambu* [21], which we also ran on AnVIL, with a novel discovery rate (NDR) threshold of 0.244. We used `splicelogic` to detect splicing events between the annotated and novel transcripts and summarized them by type, detecting 790 events in total after collapsing redundant exon events, most frequently ATSS (262, 33.2%) and ATES (219, 27.7%). Of the remaining 309 internal splicing events, the most frequent was SE (72, 23.3%).

## Conclusion

`splicelogic` turns any two sets of transcripts into discrete splicing events. Because it operates on whole transcript structures, these events are derived with full isoform context. splicelogic provides a modular, efficient, and flexible framework for identifying and interpreting splicing events while integrating with existing workflows.

## Supporting information

RBP workflow

Novel transcripts workflow

## Code availability

The splicelogic package has been part of Bioconductor since release 3.23, and is also available at https://github.com/thelovelab/splicelogic. The code used to generate the simulation and analyze the datasets can be found at https://github.com/beamimc/splicelogic-paper, archived at doi:10.5281/zenodo.23787440.

## Acknowledgments

We thank Justin Landis, Rachel Sharp, Emma Jones, Dafni Glinos, Megan Schertzer, Gloria Sheynkman, and Kateřina Matějková for helpful discussions and feedback.

## Funding

This work was supported by the National Human Genome Research Institute [R01-HG009937] and by the Wellcome Trust [313919/Z/24/Z] as part of the Essential Open Source Software for Science (EOSS) program.

## Supplementary Methods

### Matching of flanking exons

The functions `find_se()` and `find_mxe()` share the same vectorized first steps. Exons are split into the “positive” and “negative” sets by the sign of their estimate. Candidate exons are then selected with a single directed overlap count, within each gene. For each candidate, the two exons flanking it in its own transcript are matched, in one batched overlap search, against the “positive” exons of the same gene. A “positive” transcript skips the candidate when the two flanks match consecutive exons in it; for MXE, the flanks must match exons one apart, and the exon between them must not overlap the candidate. In both cases, the matched exons must not extend into the candidate. `find_ie()` similarly runs `find_se()` with the roles of the “positive” and “negative” sets swapped. The type argument sets what counts as a match between a flanking exon and its equivalent exon in the other set:

- `“over”` (default): any overlap, with no coordinate in common required. This is the most permissive setting.
- `“boundary”` : the two exons overlap *and* share the inner boundary of the flank, the splice site facing the candidate: the donor of the upstream flank and the acceptor of the downstream one. The outer boundaries may differ.
- `“identical”` : the two exons are identical (same start and same end). This is the strictest setting.

### Benchmarking of event detection functions

Simulated data were generated with `create_mock_data()`, giving each gene one “negative” transcript and one or more “positive” transcripts, each with 15 exons. Skipped-exon events were introduced with `generate_se()` in a fraction of genes set by the event ratio, with a maximum of 1 event per gene. From a baseline of 1,000 genes, an event ratio of 1, and one “positive” transcript per gene, we varied one parameter at a time: the number of genes (500, 1,000, 2,000, 5,000), the event ratio (0.1, 0.5, 1), and the number of “positive” transcripts per gene (1 to 10). Each setting was timed with *bench* over 200 iterations on a MacBook Air with an Apple M4 chip and 16 GB of memory (macOS 15.6.1). Figure 1a shows the mean ±s.e. runtime of `find_se()` with the default `type, “over”`. We also measured the runtime of `find_se()` for each type setting, and of the other event finders, varying one parameter at a time from the same baseline.

### Comparison of exon translations with Biosurfer

Exons in phase were translated directly. For exons out of phase, the one or two nucleotides of the split codon at each boundary were first trimmed, and the split-codon residue was removed from the *Biosurfer* sequence before comparing. We did not compare amino acid sequences for retained introns, because they often introduce a premature stop codon or frameshift.

### Intronic sequence flanking skipped exons

Skipped exons (SE) were obtained with `find_se()` on transcripts with significant DTU (*satuRn*, FDR < 0.1) between PTBP1 knockdown and control samples in the GTEx v9 ONT data of Glinos *et al*. [18]. Transcript abundance was quantified with *oarfish* against the GENCODE v48 transcripts. As a comparison set, we sampled an equal number of random non-skipped exons (non-SE) from transcripts without evidence of DTU (*satuRn, p* > 0.5). For each exon, we extracted the 100 bp of intronic sequence upstream and downstream with `get_seq()` and computed the nucleotide fraction in 10-bp overlapping windows sliding by 5 bp. Figure 1b shows the mean nucleotide fraction per window, with ribbons showing ±s.e.

### Collapsing events by exon and event type

Each `find_*()` call returns one row per exon and paired transcript. Because an exon can participate in an event against more than one partner transcript, the same range can appear on several rows: one for each transcript of the compared set with which it forms the same type of event. Keeping these rows is useful when both transcripts involved in each event are needed. However, for counting events, for example to summarize the total number of events by type, the rows should first be collapsed to one call per exon and event type, with the partner transcript IDs concatenated (Supplementary Code 1). The column n gives the number of transcript pairs behind each call.

**Supplementary code 1:**
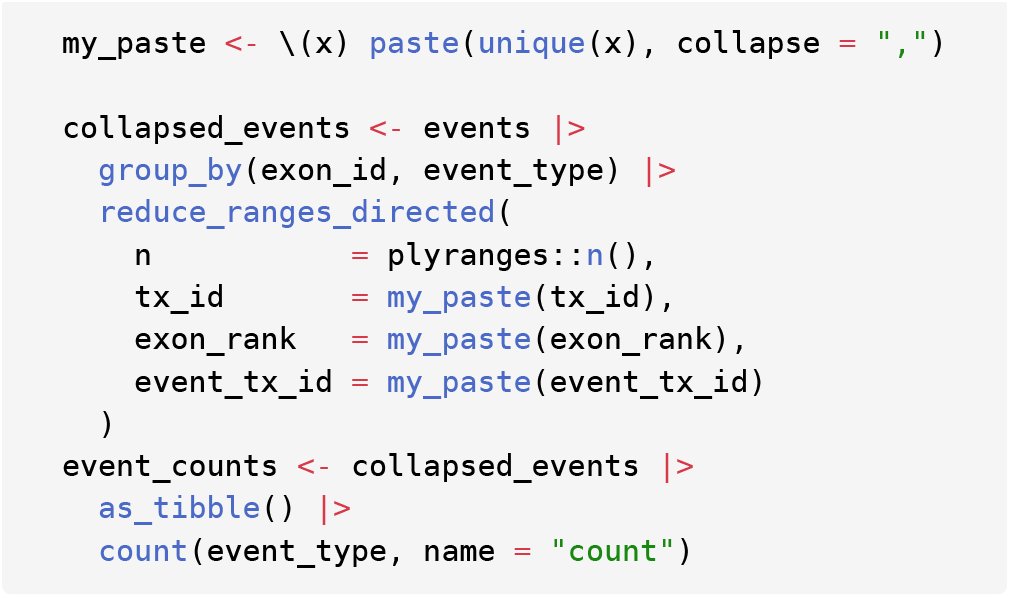
Collapsing and summarizing splicing events. This code makes use of *plyranges* and *dplyr* to manipulate genomic ranges following `splicelogic` event detection functions.

### Piping with splicelogic and plyranges

The base R pipe |> and the *magrittr* pipe %>% can usually be used interchangeably with splicelogic and *plyranges* functions. The exception is when the piped object must be referenced inside an argument on the right-hand side, as in mutate(`intron_seq = get_seq(., “hg38”`)) in Code 1. There, %>% is needed, because its. placeholder stands for the piped object.

### Software versions

R analyses used R 4.6 and Bioconductor 3.23, with `splicelogic` 1.1.4. Transcripts were quantified with *oarfish* 0.9.4 and novel isoforms detected with *bambu* 3.12.1 (R 4.5 and Bioconductor 3.22), both on AnVIL. *Biosurfer* output was based on GENCODE v42, and all other analyses used GENCODE v48.

### GTEx data

The Genotype-Tissue Expression (GTEx) Project was supported by the Common Fund of the Office of the Director of the National Institutes of Health, and by NCI, NHGRI, NHLBI, NIDA, NIMH, and NINDS. The data used for the analyses described in this manuscript were obtained from dbGaP accession number phs000424.v11.p2 in May 2026.

## Notes

### Competing Interest Statement

The authors have declared no competing interest.

