## Supplementary material for "Deducing splicing events from transcript sets with splicelogic": RBP workflow: RBP_workflow.html


### RBP workflow

```
library(SummarizedExperiment)
library(GenomicFeatures)
library(dplyr)
library(ggplot2)
library(satuRn)
library(limma)
library(tibble)
library(here)
# splicelogic devel, loaded from source: clone the repo next to this one
#   git clone -b devel https://github.com/thelovelab/splicelogic.git ../splicelogic
# then load it with devtools (install.packages("devtools") if needed)
devtools::load_all(here("../splicelogic"))
```

```
# dir <- system.file("extdata", package = "splicelogic")
# se <- readRDS(file.path(dir, "GTEx_v9_ONT_SummarizedExperiment.rds"))
se <- readRDS(here("workflows/data/GTEx_v9_ONT_SummarizedExperiment.rds"))
dim(se)
```

```
## [1] 385669     11
```

```
keep <- rowSums(assay(se) >= 10) >= 6
se <- se[keep, ]
dim(se)
```

```
## [1] 18985    11
```

```
# filters: 385669 -> 18985 transcripts

library(DESeq2)
library(tximeta)
colsums <- colSums(assay(se, "counts"))
cpm <- t(t(assay(se, "counts")) / colsums * 1e6)
assay(se, "abundance") <- cpm

# dummy lengths... no correction for length
assay(se, "length", withDimnames=FALSE) <- matrix(1000, nrow=nrow(se), ncol=ncol(se))
```

How many samples each donor contributes to each condition, before
`ctrl` / `exp` are relabelled WT / KD below.

```
samples_per_donor <- as.data.frame(colData(se)) |>
  dplyr::count(donor, condition) |>
  tidyr::pivot_wider(names_from = condition, values_from = n, values_fill = 0) |>
  dplyr::mutate(total = ctrl + exp)

# with a total row
dplyr::bind_rows(
  samples_per_donor,
  dplyr::summarise(samples_per_donor, donor = "total",
                   dplyr::across(c(ctrl, exp, total), sum))
)
```

```
## # A tibble: 5 × 4
##   donor  ctrl   exp total
##   <chr> <int> <int> <int>
## 1 QEG5      1     1     2
## 2 QV44      2     1     3
## 3 S4Z8      1     2     3
## 4 S95S      1     2     3
## 5 total     5     6    11
```

How the object was built, from `metadata(se)`. tximeta
records its own version, the quantifier and the transcriptome.
`quantInfo` holds oarfish’s `meta_info.json`
settings, which in this run do not include the oarfish version (unlike
salmon’s `salmon_version`), so that has to come from the
original run.

```
md <- metadata(se)
tibble::tibble(
  field = c("tximeta version", "quantifier", "transcriptome", "genome",
            "import time"),
  value = c(as.character(md$tximetaInfo$version), md$tximetaInfo$type,
            md$txomeInfo$index, md$txomeInfo$genome,
            format(md$tximetaInfo$importTime))
)
```

```
## # A tibble: 5 × 2
##   field           value              
##   <chr>           <chr>              
## 1 tximeta version 1.30.0             
## 2 quantifier      oarfish            
## 3 transcriptome   gencode.v48        
## 4 genome          GRCh38             
## 5 import time     2026-05-08 11:11:52
```

```
# oarfish settings, identical across the 11 samples
unique(md$quantInfo$prob_model)
```

```
## [1] "no_coverage"
```

```
unique(md$quantInfo$alignment_source)
```

```
## [1] "from_raw_reads"
```

#### DTU testing with satuRn

We compare the experiment condition (`exp`, RBP knockdown)
against control (`ctrl`) using satuRn.

```
colData(se)$condition <- dplyr::case_when(
    colData(se)$condition == "ctrl" ~ "WT",
    colData(se)$condition == "exp" ~ "KD"
)
# KD as reference level
colData(se)$condition <- factor(colData(se)$condition, levels = c("KD", "WT"))

# build design from the factor directly (matches satuRn vignette convention)
condition <- colData(se)$condition
donor <- factor(colData(se)$donor)
design <- model.matrix(~ 0 + condition + donor)

L <- limma::makeContrasts(
  WT_vs_KD = conditionWT - conditionKD,
  levels = design
)
L
```

```
##              Contrasts
## Levels        WT_vs_KD
##   conditionKD       -1
##   conditionWT        1
##   donorQV44          0
##   donorS4Z8          0
##   donorS95S          0
```

```
rowData(se)$isoform_id <- rowData(se)$tx_name
# flatten CharacterList gene_id to plain character
rowData(se)$gene_id <- vapply(rowData(se)$gene_id, `[`, character(1), 1)

se <- se[
  duplicated(rowData(se)$gene_id) |
    duplicated(rowData(se)$gene_id, fromLast = TRUE),
]
# 18985 transcripts -> 11874 transcripts after removing single-isoform genes
se <- satuRn::fitDTU(se, formula = ~0 + condition + donor)
```

```
se <- satuRn::testDTU(
  object = se,
  contrasts = L,
  diagplot1 = TRUE,
  diagplot2 = TRUE
)
```

```
gene_ids <- rowData(se)$gene_id

# every tested transcript; `sig_res` below is the significant subset, and the
# non-significant rows are the background pool for slSampleNonEventExons()
dtu_res <- rowData(se)[["fitDTUResult_WT_vs_KD"]] |>
  as.data.frame() |>
  tibble::rownames_to_column("tx_name") |>
  tibble::as_tibble() |>
  dplyr::mutate(gene_id = gene_ids) |>
  dplyr::select(tx_name, gene_id, estimates, pval, regular_FDR) |>
  dplyr::arrange(pval)

sig_res <- dtu_res |> dplyr::filter(regular_FDR < 0.1)

sig_res <- sig_res |> mutate( tx_id = tx_name)
sig_res
```

```
## # A tibble: 1,987 × 6
##    tx_name            gene_id            estimates     pval regular_FDR tx_id   
##    <chr>              <chr>                  <dbl>    <dbl>       <dbl> <chr>   
##  1 ENST00000378292.9  ENSG00000198467.16      2.98 1.00e-14    1.19e-10 ENST000…
##  2 ENST00000355572.6  ENSG00000196923.14     -3.01 5.09e-12    3.02e- 8 ENST000…
##  3 ENST00000355841.7  ENSG00000196923.14      3.63 7.77e-12    3.08e- 8 ENST000…
##  4 ENST00000403994.9  ENSG00000140416.26     -3.92 1.66e-11    4.94e- 8 ENST000…
##  5 ENST00000311785.11 ENSG00000138674.17      1.84 2.18e-11    5.17e- 8 ENST000…
##  6 ENST00000358278.7  ENSG00000140416.26      2.82 3.85e-11    7.63e- 8 ENST000…
##  7 ENST00000292807.9  ENSG00000161203.14     -2.68 6.14e-11    8.47e- 8 ENST000…
##  8 ENST00000382456.7  ENSG00000161203.14      2.68 6.14e-11    8.47e- 8 ENST000…
##  9 ENST00000613865.5  ENSG00000138326.21      1.53 6.42e-11    8.47e- 8 ENST000…
## 10 ENST00000645482.3  ENSG00000198467.16     -3.50 7.22e-11    8.58e- 8 ENST000…
## # ℹ 1,977 more rows
```

How many transcripts and genes are left after the FDR < 0.1 cut,
split by the sign of the estimate (> 0 = higher usage in WT, < 0 =
higher usage in KD). A gene counts as up (or down) when at least one of
its significant transcripts is; a gene with both is one where
splicelogic has a pair to compare.

```
sig_genes <- sig_res |>
  dplyr::group_by(gene_id) |>
  dplyr::summarise(n_up = sum(estimates > 0), n_down = sum(estimates < 0),
                   .groups = "drop")

tibble::tribble(
  ~level,        ~set,                                 ~n,
  "transcripts", "tested",                             nrow(dtu_res),
  "transcripts", "FDR < 0.1",                          nrow(sig_res),
  "transcripts", "FDR < 0.1, estimate > 0 (up, WT)",   sum(sig_res$estimates > 0),
  "transcripts", "FDR < 0.1, estimate < 0 (down, KD)", sum(sig_res$estimates < 0),
  "genes",       "tested",                             dplyr::n_distinct(dtu_res$gene_id),
  "genes",       "with >= 1 FDR < 0.1 transcript",     nrow(sig_genes),
  "genes",       "with >= 1 up transcript",            sum(sig_genes$n_up > 0),
  "genes",       "with >= 1 down transcript",          sum(sig_genes$n_down > 0),
  "genes",       "with >= 1 up and >= 1 down",         sum(sig_genes$n_up > 0 & sig_genes$n_down > 0)
) |>
  dplyr::mutate(pct = sprintf("%.1f%%", 100 * n / n[set == "tested"]),
                .by = level)
```

```
## # A tibble: 9 × 4
##   level       set                                    n pct   
##   <chr>       <chr>                              <int> <chr> 
## 1 transcripts tested                             11874 100.0%
## 2 transcripts FDR < 0.1                           1987 16.7% 
## 3 transcripts FDR < 0.1, estimate > 0 (up, WT)     953 8.0%  
## 4 transcripts FDR < 0.1, estimate < 0 (down, KD)  1034 8.7%  
## 5 genes       tested                              4369 100.0%
## 6 genes       with >= 1 FDR < 0.1 transcript      1023 23.4% 
## 7 genes       with >= 1 up transcript              849 19.4% 
## 8 genes       with >= 1 down transcript            892 20.4% 
## 9 genes       with >= 1 up and >= 1 down           718 16.4%
```

The same genes split into three groups by the direction of their
significant transcripts, with how many transcripts each group holds.
Only the “up and down” genes can give events; the other two are dropped
by the filter below.

```
gene_classes <- sig_genes |>
  dplyr::mutate(class = dplyr::case_when(
    n_up > 0 & n_down > 0 ~ "up and down",
    n_up > 0              ~ "up only",
    TRUE                  ~ "down only"
  )) |>
  dplyr::group_by(class = factor(class, levels = c("up only", "down only",
                                                   "up and down"))) |>
  dplyr::summarise(genes = dplyr::n(),
                   transcripts = sum(n_up + n_down),
                   up_tx = sum(n_up),
                   down_tx = sum(n_down)) |>
  dplyr::mutate(class = as.character(class))

# with a total row, which matches the FDR < 0.1 counts in the table above
dplyr::bind_rows(
  gene_classes,
  dplyr::summarise(gene_classes, class = "total",
                   dplyr::across(c(genes, transcripts, up_tx, down_tx), sum))
)
```

```
## # A tibble: 4 × 5
##   class       genes transcripts up_tx down_tx
##   <chr>       <int>       <int> <int>   <int>
## 1 up only       131         141   141       0
## 2 down only     174         181     0     181
## 3 up and down   718        1665   812     853
## 4 total        1023        1987   953    1034
```

```
# the significant transcripts, filtered to those with a partner in the same gene:
# keep genes with >= 1 up and >= 1 down transcript (from `sig_genes` above).
# Events always pair an up with a down transcript of the same gene, so this
# only drops transcripts that could never be in an event; the events found
# below are identical with or without it.
paired_genes <- sig_genes$gene_id[sig_genes$n_up > 0 & sig_genes$n_down > 0]
sig_res <- sig_res |> dplyr::filter(gene_id %in% paired_genes)

c(transcripts = nrow(sig_res), genes = dplyr::n_distinct(sig_res$gene_id))
```

```
## transcripts       genes 
##        1665         718
```

```
# save sig_res for later use
saveRDS(sig_res, here("workflows/data/sig_res.rds"))
```

```
# load the saved DTU results
sig_res <- readRDS(here("workflows/data/sig_res.rds"))
sig_res
```

```
## # A tibble: 1,665 × 6
##    tx_name            gene_id            estimates     pval regular_FDR tx_id   
##    <chr>              <chr>                  <dbl>    <dbl>       <dbl> <chr>   
##  1 ENST00000378292.9  ENSG00000198467.16      2.98 1.00e-14    1.19e-10 ENST000…
##  2 ENST00000355572.6  ENSG00000196923.14     -3.01 5.09e-12    3.02e- 8 ENST000…
##  3 ENST00000355841.7  ENSG00000196923.14      3.63 7.77e-12    3.08e- 8 ENST000…
##  4 ENST00000403994.9  ENSG00000140416.26     -3.92 1.66e-11    4.94e- 8 ENST000…
##  5 ENST00000311785.11 ENSG00000138674.17      1.84 2.18e-11    5.17e- 8 ENST000…
##  6 ENST00000358278.7  ENSG00000140416.26      2.82 3.85e-11    7.63e- 8 ENST000…
##  7 ENST00000292807.9  ENSG00000161203.14     -2.68 6.14e-11    8.47e- 8 ENST000…
##  8 ENST00000382456.7  ENSG00000161203.14      2.68 6.14e-11    8.47e- 8 ENST000…
##  9 ENST00000613865.5  ENSG00000138326.21      1.53 6.42e-11    8.47e- 8 ENST000…
## 10 ENST00000645482.3  ENSG00000198467.16     -3.50 7.22e-11    8.58e- 8 ENST000…
## # ℹ 1,655 more rows
```

```
library(txdbmaker)
# load TxDb for GRanges conversion
# download once (warning: ~50MB compressed)
wget https://ftp.ebi.ac.uk/pub/databases/gencode/Gencode_human/release_48/gencode.v48.annotation.gtf.gz

txdb <- txdbmaker::makeTxDbFromGFF(
  "/Users/beacm/Documents/splicelogic-paper/workflows/gencode.v48.annotation.gtf.gz",
  format = "gtf"
)
# save to not rebuild every time
AnnotationDbi::saveDb(txdb, here("workflows/data/gencode.v48.sqlite"))

# check how many transcripts in se are in txdb
rowData(se)$tx_name %in% AnnotationDbi::keys(txdb, "TXNAME") |> sum() / length(rowData(se)$tx_name)
```

#### Splice event detection with splicelogic

```
# build exon GRangesList and rename from internal TXID to TXNAME
txdb <- AnnotationDbi::loadDb(here("workflows/data/gencode.v48.sqlite"))
exons <- exonsBy(txdb, by = "tx")
txps <- AnnotationDbi::select(txdb, keys(txdb, "TXID"), c("TXNAME", "GENEID"), "TXID") |>
  tibble::as_tibble() |>
  dplyr::mutate(TXID = as.character(TXID))
```

```
## 'select()' returned 1:1 mapping between keys and columns
```

```
# rename exons to match tx_name in se, matching by TXID
all.equal(names(exons), txps$TXID)
```

```
## [1] TRUE
```

```
names(exons) <- txps$TXNAME[match(names(exons), txps$TXID)]
saveRDS(exons, here("workflows/data/glinos_exons.rds"))
```

```
# load from saved sqlite
txdb <- AnnotationDbi::loadDb(here("workflows/data/gencode.v48.sqlite"))
# load exons from saved rds
exons <- readRDS(here("workflows/data/glinos_exons.rds"))
```

```
# prepare exons for splicelogic
sig_exons <-  prepare_exons(
  txdb, sig_res, coef_col = "estimates", verbose = TRUE
)
```

```
## Extracting exons from TxDb...
```

```
## Mapping transcript IDs...
```

```
## Merging DTU results onto exons...
```

```
## Done. Returned 18990 exon ranges from 1665 unique transcripts.
```

```
#save sl_exons for later use
saveRDS(sig_exons, here("workflows/data/sl_exons_glinos_0.1.rds"))
```

```
#find se
se_events <- sig_exons |> preprocess(coef_col = "estimates")|> find_se()

#find all events
events <- sig_exons |> preprocess(coef_col = "estimates")|> find_all_events()
```

```
## Calculating skipped exon events...
```

```
## Calculating included exon events...
```

```
## Calculating mutually exclusive exon events...
```

```
## Calculating retained intron events...
```

```
## Calculating spliced intron events...
```

```
## Calculating alternative 5' splice site events...
```

```
## Calculating alternative 3' splice site events...
```

```
## Calculating alternative transcription start site events...
```

```
## Calculating alternative transcription end site events...
```

```
## Done! 2868 events detected.
```

```
# define a helper function
my_paste <- \(x) paste(unique(x), collapse = ",")
# rows are collapsed when they share the same exon and the same event type. The
# exon is identified by `exon_key` (gene, chromosome, start, end and strand),
# built as in the vignette's recipe for an exon id, rather than by `exon_id`:
# the annotation gives the same exon more than one id when it is coding in one
# transcript and non-coding in another
collapsed_events <- events |>
    dplyr::mutate(exon_key = paste(gene_id, seqnames, start, end, strand, sep = ":")) |>
    dplyr::group_by(exon_key, event_type) |>
    plyranges::reduce_ranges_directed(
        n = plyranges::n(),
        exon_id = my_paste(exon_id),
        tx_id = my_paste(tx_id),
        exon_rank = my_paste(exon_rank),
        event_tx_id = my_paste(event_tx_id),
  )
```

```
event_counts <- events |>
    as_tibble() |>
    dplyr::group_by(event_type) |>
    dplyr::summarise(count = dplyr::n())

collapsed_event_counts <- collapsed_events |>
    as_tibble() |>
    dplyr::group_by(event_type) |>
    dplyr::summarise(count = dplyr::n())

# combine both count tables into one, labelled by source
counts <- dplyr::bind_rows(
  "All events" = event_counts,
  "Collapsed events" = collapsed_event_counts,
  .id = "set"
)

# event types from most to fewest events
counts$event_type <- factor(
  counts$event_type,
  levels = event_counts$event_type[order(-event_counts$count)]
)

ggplot(counts, aes(x = event_type, y = count, fill = set)) +
  geom_col(position = position_dodge(width = 0.8)) +
  scale_fill_manual(values = c(
    "All events" = "#B0CFFF",       # light blue
    "Collapsed events" = "#3A86FF"  # darker blue
  )) +
  labs(title = "Splice event counts",
  x = "Event Type", y = "Count", fill = NULL) +
  theme_bw()
```

```
# use splicelogic's built-in plotting function to visualize events for one gene,
# picked at random among the genes with events (the seed keeps the same pick on
# every run, as long as the event set does not change)
set.seed(14)
gene_of_interest <- sample(unique(events$gene_id), 1)
# filter events for the gene of interest
events_for_gene <- events[events$gene_id == gene_of_interest]

# visualize the events for the gene of interest
splicelogic::plot_transcripts(events_for_gene, sig_exons, rescale_introns = TRUE) +
  ggtitle(paste("Splice events for gene:", gene_of_interest))
```

```
# make a plot for each gene with events and save to files
# plot_transcripts_by_gene returns a list of ggplot objects, one for each gene
plots <- plot_transcripts_by_gene(events, sig_exons)
# save each plot to the same pdf file - takes a while: -- 1725 genes 
pdf(here("workflows/figures/splice_events_by_gene.pdf"), width = 8, height = 6)
for (gene in names(plots)) {
    print(plots[[gene]])
}
dev.off()
```

#### Collapse redundant events

`find_se()` returns an exon once per transcript pair it is
part of: an exon in two down transcripts and skipped by two up ones
comes back four times. The event counts above already collapse these
(`collapsed_events`). For the sequence analyses below we also
keep one row per exon, so that an exon shared by many transcripts does
not weigh more in the profiles and logos, and the background set is
sized by exons rather than pairs. `n` records how many pairs
each exon came from. The collapse is the same as in
`collapse_redundant_events`, applied to the SE events, and
like it collapses rows only for the same exon (gene, chromosome, start,
end, strand) and the same event type.

```
# TRUE: the sequence analyses below use one row per exon. FALSE: they use every
# SE event, one row per transcript pair, as before the collapse, to compare
collapse_se <- TRUE

se_collapsed <- se_events |>
    dplyr::mutate(exon_key = paste(gene_id, seqnames, start, end, strand, sep = ":")) |>
    dplyr::group_by(exon_key, event_type) |>
    plyranges::reduce_ranges_directed(
        n = plyranges::n(),
        gene_id = my_paste(gene_id),
        exon_id = my_paste(exon_id),
        tx_id = my_paste(tx_id),
        exon_rank = my_paste(exon_rank),
        event_tx_id = my_paste(event_tx_id),
  )

data.frame(
  set    = c("SE"),
  events = c(length(se_events)),
  exons  = c(length(se_collapsed)),
  max_n  = c(max(se_collapsed$n))
)
```

```
##   set events exons max_n
## 1  SE    306   228    12
```

```
# the set the sequence analyses below use
se_exons <- if (collapse_se) se_collapsed else se_events
```

##### SE exons per gene

How the SE events spread across genes: the number of distinct SE
exons in each gene. `se_collapsed` already has one row per
exon and gene, with `n` the events it came from, so the
counts come straight from it.

```
se_per_gene <- se_collapsed |>
  dplyr::group_by(gene_id) |>
  dplyr::summarise(n_events = sum(n),
                   n_exons  = plyranges::n())

data.frame(
  se_events = length(se_events),
  se_exons  = length(se_collapsed),
  se_genes  = nrow(se_per_gene)
)
```

```
##   se_events se_exons se_genes
## 1       306      228      190
```

```
# ggplot needs a data.frame, so the small per-gene summary is converted here
as.data.frame(se_per_gene) |>
  dplyr::count(n_exons) |>
  ggplot(aes(x = factor(n_exons), y = n)) +
  geom_col(fill = "#3A86FF", width = 0.6) +
  geom_text(aes(label = n), vjust = -0.4, size = 3.5, colour = "grey20") +
  scale_y_continuous(expand = expansion(mult = c(0, 0.1))) +
  labs(title = "Distinct SE exons per gene",
       subtitle = sprintf("%d SE events, %d distinct exons, %d genes",
                          length(se_events), length(se_collapsed),
                          nrow(se_per_gene)),
       x = "Distinct SE exons in the gene", y = "Genes") +
  theme_bw()
```

##### GO enrichment of SE genes

Which biological processes are the genes with an SE event enriched
for? The test set is every gene with at least one SE event; the
background (universe) is every gene tested for DTU, since only those
could give an event. Testing against the whole genome instead would
mostly pick up which genes are expressed with several isoforms in this
tissue. GO biological process terms, over-representation test with
Benjamini-Hochberg adjustment, and
`clusterProfiler::simplify()` to merge near-identical
terms.

```
se_genes  <- unique(sub("\\..*", "", as.character(se_events$gene_id)))
universe  <- unique(sub("\\..*", "", dtu_res$gene_id))
c(se_genes = length(se_genes), universe = length(universe))
```

```
## se_genes universe 
##      190     4369
```

```
go_se <- clusterProfiler::enrichGO(
  gene          = se_genes,
  universe      = universe,
  OrgDb         = org.Hs.eg.db::org.Hs.eg.db,
  keyType       = "ENSEMBL",
  ont           = "BP",
  pAdjustMethod = "BH",
  pvalueCutoff  = 0.05,
  qvalueCutoff  = 0.2,
  readable      = TRUE   # gene symbols in the geneID column
)
```

```
## Warning in bitr(gene, fromType = fromType, toType = "ENTREZID", OrgDb = OrgDb):
## 1.14% of input gene IDs are fail to map...
```

```
# merge redundant terms (semantic similarity > 0.7), keeping the most
# significant of each group
go_se <- clusterProfiler::simplify(go_se, cutoff = 0.7, by = "p.adjust",
                                   select_fun = min)
```

```
## Warning: semData is not provided. It will be calculated automatically.
```

```
go_se_df <- as.data.frame(go_se)
nrow(go_se_df)
```

```
## [1] 19
```

```
go_se_df |>
  dplyr::select(Description, GeneRatio, BgRatio, p.adjust, geneID) |>
  head(20)
```

```
##                                                               Description
## GO:1990778                         protein localization to cell periphery
## GO:0007155                                                  cell adhesion
## GO:0051668                                   localization within membrane
## GO:0007015                                    actin filament organization
## GO:0032880                             regulation of protein localization
## GO:0000902                                             cell morphogenesis
## GO:0060341                            regulation of cellular localization
## GO:0044319                              wound healing, spreading of cells
## GO:0016192                                     vesicle-mediated transport
## GO:0032231                   regulation of actin filament bundle assembly
## GO:1903077 negative regulation of protein localization to plasma membrane
## GO:0030155                                    regulation of cell adhesion
## GO:0030029                                   actin filament-based process
## GO:0042060                                                  wound healing
## GO:1903829                    positive regulation of protein localization
## GO:0015031                                              protein transport
## GO:0045184                          establishment of protein localization
## GO:0060627                       regulation of vesicle-mediated transport
## GO:0097435                              supramolecular fiber organization
##            GeneRatio  BgRatio    p.adjust
## GO:1990778    19/180 109/3941 0.000586719
## GO:0007155    36/180 354/3941 0.001611742
## GO:0051668    26/180 218/3941 0.001628784
## GO:0007015    19/180 132/3941 0.001998026
## GO:0032880    28/180 265/3941 0.004477878
## GO:0000902    23/180 206/3941 0.009157586
## GO:0060341    20/180 173/3941 0.014048941
## GO:0044319     6/180  19/3941 0.018529719
## GO:0016192    36/180 433/3941 0.019633980
## GO:0032231     8/180  37/3941 0.019633980
## GO:1903077     5/180  14/3941 0.022596137
## GO:0030155    21/180 203/3941 0.023934291
## GO:0030029    22/180 220/3941 0.024780247
## GO:0042060    14/180 111/3941 0.030248338
## GO:1903829    17/180 153/3941 0.032152667
## GO:0015031    34/180 421/3941 0.032152667
## GO:0045184    38/180 491/3941 0.032152667
## GO:0060627    15/180 129/3941 0.034644712
## GO:0097435    22/180 233/3941 0.037687262
##                                                                                                                                                                                                                                         geneID
## GO:1990778                                                                                                              MACF1/ATP2B4/TTC7A/CCDC88A/GOLGA4/ATP2C1/ATP1B3/AP2M1/LRRC15/AP3B1/ILK/PPFIA1/PICALM/VTI1B/MYO5A/CLTC/IFT20/EXOC7/FLNA
## GO:0007155                      MACF1/NEXN/CD46/MAP4K4/PKP4/ADAM17/CLASP1/DPP4/FN1/FBLN2/ATP2C1/RNF7/BCL6/VCAN/AP3B1/PPIA/ASS1/CD151/ILK/CTNND1/PPFIA1/PPFIBP1/PPP1R12A/PXN/POSTN/FERMT2/FBLN5/ATXN3/TPM1/CDH11/PRKAR1A/PLAUR/GNAS/NF2/XG/FLNA
## GO:0051668                                                                   MACF1/ATP2B4/SRP9/TTC7A/CCDC88A/GOLGA4/ATP2C1/ATP1B3/FXR1/AP2M1/LRRC15/AP3B1/SIL1/PPFIA1/PICALM/TIMM8B/VPS29/FERMT2/VTI1B/SRP14/MYO5A/CLTC/IFT20/EXOC7/MTCL1/FLNA
## GO:0007015                                                                                                                           CAPZB/TPM3/ENAH/CCDC88A/CLASP1/DBN1/CALD1/SPTAN1/ABI1/SVIL/ILK/PPFIA1/PXN/FERMT2/TPM1/MYO5A/TPM4/NF2/FLNA
## GO:0032880                                                        ATP2B4/CCDC88A/ATP2C1/AP2M1/SEC13/PDCD10/PARL/LRRC15/SEC31A/ABLIM3/PPIA/CCT6A/ASPH/SEC24C/PPP3CB/CTNND1/PPFIA1/PICALM/OS9/MARK3/FERMT2/VTI1B/CLTC/MTCL1/GNAS/NSFL1C/NF2/FLNA
## GO:0000902                                                                                            MACF1/NEXN/MOV10/CDC42SE1/ENAH/MAP4K4/ADAM17/FN1/GOLGA4/DBN1/ENPP2/ABI1/PPP3CB/ILK/APLP2/PICALM/PPP1R12A/FERMT2/TPM1/CDH11/TPM4/NF2/FLNA
## GO:0060341                                                                                                             ATP2B4/ATP2C1/AP2M1/SEC13/PDCD10/LRRC15/SEC31A/ASPH/SEC24C/PPP3CB/PPFIA1/PICALM/OS9/ISCU/VTI1B/CLTC/MTCL1/GNAS/NF2/FLNA
## GO:0044319                                                                                                                                                                                              ADAM17/CLASP1/PDCD10/CD151/FERMT2/FLNA
## GO:0016192 MACF1/PPT1/YIPF1/CDC42SE1/CLASP1/DPP4/GOLGA4/ATP2C1/AP2M1/SEC13/OCIAD2/SEC31A/AP3B1/SMAP1/TMED4/PPP3CC/ENPP2/CLTA/GOLGA2/FNBP1/SEC24C/PPP3CB/CD151/PICALM/GOLT1B/VPS29/TRAPPC6B/VTI1B/MYO5A/CLTC/EXOC7/TRIP10/AXL/EPS15L1/AP2S1/NF2
## GO:0032231                                                                                                                                                                                      CCDC88A/CLASP1/PPFIA1/PXN/FERMT2/TPM1/NF2/FLNA
## GO:1903077                                                                                                                                                                                                     AP2M1/LRRC15/PPFIA1/PICALM/CLTC
## GO:0030155                                                                                                             MACF1/CD46/MAP4K4/PKP4/CLASP1/DPP4/FN1/FBLN2/RNF7/BCL6/AP3B1/ASS1/ILK/PPP1R12A/FERMT2/ATXN3/TPM1/PRKAR1A/PLAUR/NF2/FLNA
## GO:0030029                                                                                                       CAPZB/TPM3/ENAH/CCDC88A/CLASP1/ATP2C1/DBN1/CALD1/SPTAN1/ABI1/SVIL/ILK/PPFIA1/PXN/FERMT2/ATXN3/TPM1/MYO5A/TRIP10/TPM4/NF2/FLNA
## GO:0042060                                                                                                                                                       MACF1/ADAM17/CLASP1/FN1/PDCD10/PPIA/CD151/ILK/FERMT2/TPM1/AXL/PLAUR/GNAS/FLNA
## GO:1903829                                                                                                                          ATP2B4/CCDC88A/ATP2C1/SEC13/PDCD10/SEC31A/ABLIM3/PPIA/CCT6A/ASPH/SEC24C/PPP3CB/CTNND1/FERMT2/GNAS/NF2/FLNA
## GO:0015031                        MACF1/PPT1/ATG16L1/GOLGA4/ATP2C1/AP2M1/SEC13/PDCD10/SEC31A/TNPO1/AP3B1/KIF3A/SIL1/PPIA/TMED4/ASPH/CLTA/GOLGA2/SEC24C/PPP3CB/TIMM8B/GOLT1B/OS9/VPS29/VTI1B/SRP14/MYO5A/CLTC/STRADA/EXOC7/AP2S1/HM13/GNAS/FLNA
## GO:0045184 MACF1/SRP9/PPT1/ATG16L1/GOLGA4/ATP2C1/AP2M1/SEC13/PDCD10/SEC31A/TNPO1/AP3B1/KIF3A/SIL1/PPIA/CCT6A/TMED4/ASPH/CLTA/GOLGA2/SEC24C/PPP3CB/SPCS2/TIMM8B/GOLT1B/OS9/VPS29/VTI1B/SRP14/MYO5A/CLTC/STRADA/EXOC7/MTCL1/AP2S1/HM13/GNAS/FLNA
## GO:0060627                                                                                                                                         PPT1/CLASP1/ATP2C1/AP2M1/SEC13/SEC31A/SMAP1/PPP3CC/SEC24C/PPP3CB/CD151/PICALM/AXL/AP2S1/NF2
## GO:0097435                                                                                                      CAPZB/TPM3/ENAH/CCDC88A/CLASP1/DBN1/CALD1/SPTAN1/GOLGA2/ABI1/SVIL/ILK/PPFIA1/PPFIBP1/PXN/FERMT2/FBLN5/TPM1/MYO5A/TPM4/NF2/FLNA
```

```
if (nrow(go_se_df) > 0) {
  enrichplot::dotplot(go_se, showCategory = 15) +
    labs(title = "GO biological process: genes with an SE event",
         subtitle = sprintf("%d SE genes vs %d DTU-tested genes",
                            length(intersect(se_genes, universe)), length(universe)))
}
```

#### Sequence context around the event exons, focusing on skipped exons (SE)

The analysis below uses a set of sequence-context helpers
consolidated in `helper.R` (nucleotide-composition profiles,
background exon sampling, splice-site logos and RBP motif scanning). The
splicelogic package itself only exports `get_seq()` for
sequence work, so these stay as paper-repo helpers (for now).

```
source(here("workflows/helper.R"))
```

`slEventFlankProfiles()` takes a finder’s output straight
away: it flanks each exon (strand-aware, so “upstream” is 5’ in
transcription order), slides overlapping windows across the flank, and
returns the per-window nucleotide fractions for each side.

```
se_profiles <- slEventFlankProfiles(se_exons, genome = "hg38", width = 100)

# rows = exons, cols = window_nucleotide (e.g. w1_A)
dim(se_profiles$upstream)
```

```
## [1] 228  76
```

```
dim(se_profiles$downstream)
```

```
## [1] 228  76
```

##### Background set

We want to compare the sequence context of the events with that of a
control set. For the control set (non-event exons), we sample as many
exons as there are in `se_exons` (one per SE exon, or one per
SE event with `collapse_se = FALSE`), drawn at random from
the internal exons of transcripts with no evidence of differential usage
(`pval > 0.5`).

```
non_event <- slSampleNonEventExons(
  dtu_res, exons, n = se_exons, pval_min = 0.5, seed = 1
)
```

```
non_event_profiles <- slEventFlankProfiles(
  non_event, genome = "hg38", width = 100
)
dim(non_event_profiles$upstream)
```

```
## [1] 228  76
```

The SE profiles stacked over the non-SE ones, on the same axes:

```
p_se <- slPlotFlankProfiles(se_profiles, exon_label = "SE")

# one shared nucleotide legend at the top
patchwork::wrap_plots(
  p_se,
  slPlotFlankProfiles(non_event_profiles, exon_label = "non-SE"),
  ncol = 1, guides = "collect"
) & theme(legend.position = "top")
```

```
# One text size for every poster figure, in pt at 300 dpi: tick labels, axis
# titles, legends and labels inside the plots all use it. se_composition and
# rna_map are drawn at 3.6 x 3 in and 600 dpi, half the size at twice the
# resolution, so they use poster_pt / 2 to come out the same size in pixels.
poster_pt <- 24
half_pt <- poster_pt / 2
same_text <- theme(text = element_text(size = half_pt),
                   axis.text = element_text(size = half_pt),
                   axis.title = element_text(size = half_pt),
                   legend.text = element_text(size = half_pt))

# the SE profile alone, for the poster, styled to sit side by side with the RNA
# map: the same size (3.6 x 3 in), text size, line width and legend layout.
# No plot or legend title (the poster panel's heading and caption carry them),
# but both axis titles are kept. Legend on the top, in one row.
p_se_poster <- slPlotFlankProfiles(se_profiles, exon_label = "SE",
                                   ylim = c(0, 0.5), hlines = seq(0, 0.4, by = 0.2),
                                   base_size = half_pt, linewidth = 0.8) +
  labs(title = NULL) +
  same_text +
  theme(legend.title = element_blank(),
        legend.position = "top",
        legend.direction = "horizontal",
        legend.key.width = grid::unit(0.5, "cm"),
        legend.key.height = grid::unit(7, "pt"),
        legend.box.spacing = grid::unit(2, "pt"),
        legend.margin = margin(0, 0, 0, 0),
        plot.margin = margin(2, 2, 2, 2))
dir.create(here("workflows", "figures", "poster"), showWarnings = FALSE, recursive = TRUE)
# a 3.6 x 3 in frame at 600 dpi
ggsave(here("workflows", "figures", "poster", "se_composition.png"), p_se_poster,
       width = 3.6, height = 3, dpi = 600)
```

The two sets, one panel per nucleotide (mean ± standard error), and
below them the difference between them, SE minus non-SE:

```
se_vs_non_event <- list(SE = se_profiles, "non-SE" = non_event_profiles)

p_overlay <- slPlotFlankProfilesByNt(se_vs_non_event)
p_overlay
```

```
slPlotFlankProfilesByNt(se_vs_non_event, difference = TRUE)
```

```
# poster version: same width as the RNA map (3.6 in), stacked under
# se_composition
# Large text for the poster and no axis or legend titles (the poster caption
# carries them); the nucleotide as a label to the left of each row rather than
# a title above it, to leave the 4 short rows to the data.
p_overlay_poster <- slPlotFlankProfilesByNt(se_vs_non_event,
                                            base_size = 8, linewidth = 0.7) +
  facet_wrap(~ nt, ncol = 1, strip.position = "left", scales = "free_y") +
  theme(axis.title = element_blank(),
        legend.title = element_blank(),
        strip.placement = "outside",
        strip.text.y.left = element_text(angle = 0, face = "bold", size = 10),
        panel.spacing.y = grid::unit(0.15, "lines"),
        legend.key.width = grid::unit(0.8, "cm"),
        legend.box.spacing = grid::unit(2, "pt"),
        legend.margin = margin(0, 0, 0, 0),
        plot.margin = margin(2, 2, 2, 2))
# a 3.6 in frame at 600 dpi, text and lines halved as for se_composition
ggsave(here("workflows", "figures", "poster", "se_vs_background.png"), p_overlay_poster,
       width = 3.6, height = 2.325, dpi = 600)
```

An alternative, within-transcript control: for each event, the
internal exon of the same transcript that is shared with the event
partner and closest in rank to the event exon.

```
matched <- slMatchedNonEventExons(se_events, exons)

slEventFlankProfiles(matched, genome = "hg38", width = 100) |>
  slPlotFlankProfiles(exon_label = "matched exon")
```

#### Splice-site sequence logos

The window profiles above average over 10 bp, which smooths the
splice site away. `slFlankLogoMatrices()` keeps every base
position separate and returns a position frequency matrix per side,
running `width` bases into the intron and
`exon_bases` into the exon so the boundary is inside the
matrix.

Note which site each side carries: the region
**upstream** of an exon is the end of the preceding intron
and holds the **3’ splice site** (acceptor,
`...ag | EXON`); the region **downstream** is
the start of the following intron and holds the **5’ splice
site** (donor, `EXON | gu...`).

```
se_logo <- slFlankLogoMatrices(se_exons, genome = "hg38",
                               width = 25, exon_bases = 3)

# letter height = information content, the classic logo scaling: unconstrained
# positions shrink to nothing and the consensus stands out
slPlotFlankLogo(se_logo, method = "bits")
```

The same for the background set, to see whether the event splice
sites differ from those of exons in transcripts with no differential
usage.

```
non_event_logo <- slFlankLogoMatrices(non_event, genome = "hg38",
                                      width = 25, exon_bases = 3)

slPlotFlankLogo(non_event_logo, method = "bits")
```

```
# where the two sets differ most, per position and nucleotide
delta <- se_logo$downstream - non_event_logo$downstream
round(delta[, order(-apply(abs(delta), 2, max))[1:6]], 3)
```

```
##        5     19      6     -3     13     25
## A  0.079 -0.075  0.009 -0.088  0.079 -0.044
## C  0.026  0.114  0.013  0.035 -0.013  0.075
## G -0.136  0.000  0.079  0.022 -0.070 -0.039
## U  0.031 -0.039 -0.101  0.031  0.004  0.009
```

#### PTBP1 RNA map

Colours below are taken from the package, so they match
`plot_transcripts()`: the up colour for transcripts with
higher usage in WT (estimate > 0), the down colour for higher usage
in KD, and `event_palette()` for event types.

```
# the package's own colours, read from plot_transcripts() so they stay in sync
sl_up     <- formals(plot_transcripts)$up_color       # higher usage in WT
sl_down   <- formals(plot_transcripts)$down_color     # higher usage in KD
sl_grey   <- formals(plot_transcripts)$unknown_color  # not significant / other
sl_events <- event_palette()

# keep the standard chromosomes only, so every range is in the BSgenome
standard_chr <- function(gr) {
  gr <- GenomeInfoDb::keepStandardChromosomes(gr, pruning.mode = "coarse")
  GenomeInfoDb::seqlevels(gr) <- GenomeInfoDb::seqlevelsInUse(gr)
  gr
}

# the unfiltered counts: the per-sample usage plots use every transcript of a
# gene, not only those kept for DTU
se_raw   <- readRDS(here("workflows/data/GTEx_v9_ONT_SummarizedExperiment.rds"))
raw_gid  <- sub("\\..*", "", vapply(rowData(se_raw)$gene_id, `[`, character(1), 1))
raw_cond <- ifelse(se_raw$condition == "ctrl", "WT", "KD")
raw_cnt  <- assay(se_raw, "counts")
```

SE exons are on down transcripts (higher usage in KD) and skipped in
up transcripts (higher in WT): with PTBP1 present the exon is skipped,
so these are PTBP1-induced skipping events. From PTBP1’s RNA map we
expect CU-rich motifs **upstream** of SE exons (around the
3’ splice site), relative to the background exons.

The motif is YCUY (Y = C or U), counted at every start position with
overlapping matches allowed, in `W` nt of intron on each side
of the exon. Each exon gives a 0/1 vector (1 where a motif starts),
smoothed with a centred rolling mean; the profile is the mean ± s.e.
across exons. The plot uses the same layout as the flank profiles above:
one axis running through both introns, with the exon as a grey bar
between them.

```
W <- 200
smooth_nt <- 21
motif_regex <- "(?=[CU]CU[CU])"  # lookahead, so overlapping hits all count

# 0/1 matrix, exons x positions: 1 where a motif starts. get_seq() returns the
# sequence in transcription order (minus strand reverse-complemented), and the
# flanks are strand-aware, so column 1 of "upstream" is -W and column W is -1.
flank_matrix <- function(gr, side) {
  gr <- standard_chr(gr)
  fl <- if (side == "upstream") {
    plyranges::flank_upstream(gr, width = W)
  } else {
    plyranges::flank_downstream(gr, width = W)
  }
  seqs <- as.character(get_seq(fl, "hg38", as_rna = TRUE))
  m <- matrix(0L, nrow = length(seqs), ncol = W)
  for (i in seq_along(seqs)) {
    pos <- gregexpr(motif_regex, seqs[i], perl = TRUE)[[1]]
    pos <- pos[pos > 0]
    m[i, pos] <- 1L
  }
  m
}

sets <- list(SE = se_collapsed, background = non_event)
mats <- lapply(sets, function(gr) list(
  upstream   = flank_matrix(gr, "upstream"),
  downstream = flank_matrix(gr, "downstream")
))
sapply(mats, \(m) nrow(m$upstream))  # exons per set
```

```
##         SE background 
##        228        228
```

```
# per-exon rolling mean over smooth_nt, then mean +/- s.e. across exons. As in
# slPlotFlankProfilesByNt(), both introns share one x axis: the upstream intron
# at x = 1..W (-W..-1 nt), then a gap holding the exon bar, then the
# downstream intron (+1..+W nt).
gap <- 60
profile_df <- function(m, side, set) {
  sm <- t(apply(m, 1, \(x) as.numeric(stats::filter(x, rep(1 / smooth_nt, smooth_nt), sides = 2))))
  pos <- if (side == "upstream") -W:-1 else 1:W
  tibble::tibble(
    set  = set,
    side = side,
    x    = if (side == "upstream") pos + W + 1 else pos + W + gap,
    mean = colMeans(sm, na.rm = TRUE),
    se   = apply(sm, 2, sd, na.rm = TRUE) / sqrt(nrow(sm))
  )
}

set_labels <- sprintf("%s (n = %d)", names(mats), sapply(mats, \(m) nrow(m$upstream)))
map_df <- dplyr::bind_rows(lapply(names(mats), function(s) dplyr::bind_rows(
  profile_df(mats[[s]]$upstream, "upstream", s),
  profile_df(mats[[s]]$downstream, "downstream", s)
))) |>
  dplyr::mutate(set = factor(set, levels = names(mats), labels = set_labels))

# x axis: nt from the exon every 100, the splice sites at the inner ends, and
# the exon bar in the middle (same labelling as .slFlankXAxis())
exon_x <- W + (gap + 1) / 2
bp <- seq(100, W, by = 100)
x_breaks <- c(rev(-bp) + W + 1, W, exon_x, W + gap + 1, bp + W + gap)
x_labels <- c(rev(-bp), "\n3'ss", "exon", "\n5'ss", paste0("+", bp))

# SE in its event_palette() colour, solid; background grey, dashed
map_cols <- setNames(c(sl_events[["se"]], "grey50"), set_labels)
map_lty  <- setNames(c("solid", "dashed"), set_labels)

ggplot(map_df, aes(x, mean, group = interaction(side, set))) +
  geom_ribbon(aes(ymin = mean - se, ymax = mean + se, fill = set),
              alpha = 0.25, colour = NA, na.rm = TRUE) +
  geom_line(aes(colour = set, linetype = set), linewidth = 1, na.rm = TRUE) +
  # the exon as a grey bar in the gap, and a black x-axis line under each
  # intron, broken at the exon
  .slFlankExonBar(W, gap, exon_x) +
  .slFlankXLine(W, gap) +
  coord_cartesian(clip = "off") +
  scale_colour_manual(values = map_cols) +
  scale_fill_manual(values = map_cols, guide = "none") +
  scale_linetype_manual(values = map_lty) +
  scale_x_continuous(breaks = x_breaks, labels = x_labels) +
  # from 0, with no padding below, so the x-axis line sits exactly on 0
  scale_y_continuous(limits = c(0, NA), expand = expansion(mult = c(0, 0.02))) +
  labs(title = "PTBP1 RNA map: YCUY motifs around SE exons",
       subtitle = sprintf("%d-nt rolling mean ± s.e.", smooth_nt),
       x = "Relative position in intronic sequence (nt)",
       y = "YCUY motif density per nt", colour = NULL, linetype = NULL) +
  .slFlankTheme() +
  theme(legend.key.width = grid::unit(1.2, "cm"))
```

Caveats: a 4-mer cannot start in the last 3 upstream positions
without running into the exon, so those columns are always 0; the
overlapping count gives long CU runs, such as the polypyrimidine tract,
a lot of weight; and flanks of introns shorter than `W` run
into the neighbouring exon.

#### Example SE gene: SMAP1

Two panels: isoform usage per sample, and the events
`find_all_events()` called in the gene. SMAP1 has two
isoforms that differ by one 81-nt exon (plus their 5’ and 3’ ends). The
exon-including isoform has higher usage in KD, so PTBP1 represses the
exon. To plot another gene, add it to `example_genes`.

Usage is each transcript’s share of the gene’s counts in the
unfiltered matrix. Each DTU transcript (FDR < 0.1, the ones the
events come from) gets its own colour: shades of the up colour for
higher usage in WT, shades of the down colour for higher in KD. Every
other transcript of the gene is pooled as “other”.

```
example_genes <- c(SMAP1 = "ENSG00000112305")

# KD first, then WT, numbered within condition in donor order
ord <- order(factor(raw_cond, levels = c("KD", "WT")), se_raw$donor, colnames(se_raw))
samp_lab <- character(ncol(se_raw))
samp_lab[ord] <- ave(raw_cond[ord], raw_cond[ord], FUN = \(x) paste0(x, "-", seq_along(x)))

# the package colour, then two lighter shades of it, for genes with more than
# one DTU transcript in a direction
shades <- \(col) grDevices::colorRampPalette(c(col, "white"))(5)[1:3]
pal_up   <- shades(sl_up)
pal_down <- shades(sl_down)

plot_example_gene <- function(gene, gene_id) {
  ex_events <- events[sub("\\..*", "", events$gene_id) == gene_id]
  ex_exons  <- sig_exons[sub("\\..*", "", sig_exons$gene_id) == gene_id]

  # the DTU transcripts, WT-up first, each ordered by effect size
  ex_res <- dtu_res |>
    dplyr::filter(tx_name %in% unique(ex_exons$tx_id)) |>
    dplyr::arrange(dplyr::desc(estimates > 0), dplyr::desc(abs(estimates)))
  up_tx   <- ex_res$tx_name[ex_res$estimates > 0]
  down_tx <- ex_res$tx_name[ex_res$estimates < 0]

  # legend labels: the transcript id alone
  labs_tx <- setNames(ex_res$tx_name, ex_res$tx_name)

  gene_cnt <- raw_cnt[raw_gid == gene_id, , drop = FALSE]
  iso <- ifelse(rownames(gene_cnt) %in% names(labs_tx),
                labs_tx[rownames(gene_cnt)], "other")
  iso_levels <- c(labs_tx[up_tx], labs_tx[down_tx], "other")
  iso_cols <- setNames(c(pal_up[seq_along(up_tx)], pal_down[seq_along(down_tx)], sl_grey),
                       iso_levels)

  usage_df <- rowsum(gene_cnt, iso) |>
    sweep(2, colSums(gene_cnt), "/") |>
    as.data.frame() |>
    tibble::rownames_to_column("isoform") |>
    tidyr::pivot_longer(-isoform, names_to = "sample", values_to = "usage") |>
    dplyr::mutate(
      sample  = factor(samp_lab[match(sample, colnames(se_raw))], levels = samp_lab[ord]),
      isoform = factor(isoform, levels = iso_levels),
      # "KD-3" -> condition "KD" (one shared label under its samples), "3"
      condition = factor(sub("-.*", "", sample), levels = c("KD", "WT")),
      sample_n  = factor(as.integer(sub(".*-", "", sample)))
    )

  p_usage <- ggplot(usage_df, aes(sample_n, usage, fill = isoform)) +
    geom_col(width = 0.85) +
    # one panel per condition, its name once below its sample numbers
    facet_grid(~ condition, scales = "free_x", space = "free_x", switch = "x") +
    scale_fill_manual(values = iso_cols, drop = FALSE) +
    # zoom, not scale limits: a stack can sum to 1 + rounding error, and scale
    # limits would drop that top segment as out of range
    scale_y_continuous(expand = expansion(mult = c(0, 0.02))) +
    coord_cartesian(ylim = c(0, 1)) +
    labs(title = sprintf("%s: isoform usage per sample", gene),
         x = NULL, y = "Share of gene expression", fill = NULL) +
    theme_bw() +
    theme(strip.placement = "outside",
          strip.background = element_blank(),
          panel.grid.major.x = element_blank())

  # the events splicelogic called in this gene, on the transcripts they came
  # from
  p_events <- plot_transcripts(ex_events, ex_exons, rescale_introns = TRUE) +
    labs(title = sprintf("%s: events called by find_all_events()", gene))

  list(usage = p_usage, events = p_events)
}

example_plots <- Map(plot_example_gene, names(example_genes), example_genes)
for (p in example_plots) {
  print(p$events)
  print(p$usage)
}
```

#### Session info

The R and package versions (satuRn, splicelogic, Bioconductor, …)
this report was knitted with.

```
sessionInfo()
```

```
## R version 4.6.1 (2026-06-24)
## Platform: aarch64-apple-darwin23
## Running under: macOS Sequoia 15.6.1
## 
## Matrix products: default
## BLAS:   /Library/Frameworks/R.framework/Versions/4.6/Resources/lib/libRblas.0.dylib 
## LAPACK: /Library/Frameworks/R.framework/Versions/4.6/Resources/lib/libRlapack.dylib;  LAPACK version 3.12.1
## 
## locale:
## [1] en_US.UTF-8/en_US.UTF-8/en_US.UTF-8/C/en_US.UTF-8/en_US.UTF-8
## 
## time zone: America/New_York
## tzcode source: internal
## 
## attached base packages:
## [1] stats4    stats     graphics  grDevices utils     datasets  methods  
## [8] base     
## 
## other attached packages:
##  [1] BSgenome.Hsapiens.UCSC.hg38_1.4.5 BSgenome_1.80.0                  
##  [3] rtracklayer_1.72.0                BiocIO_1.22.0                    
##  [5] Biostrings_2.80.2                 XVector_0.52.0                   
##  [7] GenomeInfoDb_1.48.0               tximeta_1.30.1                   
##  [9] DESeq2_1.52.0                     splicelogic_1.1.4                
## [11] testthat_3.3.2                    plyranges_1.32.0                 
## [13] here_1.0.2                        tibble_3.3.1                     
## [15] limma_3.68.5                      satuRn_1.20.0                    
## [17] ggplot2_4.0.3                     dplyr_1.2.1                      
## [19] GenomicFeatures_1.64.0            AnnotationDbi_1.74.0             
## [21] SummarizedExperiment_1.42.0       Biobase_2.72.0                   
## [23] GenomicRanges_1.64.0              Seqinfo_1.2.0                    
## [25] IRanges_2.46.0                    S4Vectors_0.50.3                 
## [27] BiocGenerics_0.58.1               generics_0.1.4                   
## [29] MatrixGenerics_1.24.0             matrixStats_1.5.0                
## 
## loaded via a namespace (and not attached):
##   [1] splines_4.6.1            bitops_1.1-0             ggplotify_0.1.4         
##   [4] filelock_1.0.3           polyclip_1.10-7          enrichit_0.2.7          
##   [7] XML_3.99-0.25            lifecycle_1.0.5          httr2_1.3.0             
##  [10] rprojroot_2.1.1          MASS_7.3-65              processx_3.9.0          
##  [13] lattice_0.22-9           ensembldb_2.36.1         magrittr_2.0.5          
##  [16] sass_0.4.10              rmarkdown_2.32           jquerylib_0.1.4         
##  [19] yaml_2.3.12              otel_0.2.0               ggtangle_0.1.4          
##  [22] sessioninfo_1.2.4        pkgbuild_1.4.8           pbapply_1.7-5           
##  [25] DBI_1.3.0                RColorBrewer_1.1-3       abind_1.4-8             
##  [28] pkgload_1.5.3            purrr_1.2.2              AnnotationFilter_1.36.0 
##  [31] RCurl_1.98-1.20          yulab.utils_0.2.5        tweenr_2.0.3            
##  [34] rappdirs_0.3.4           aisdk_1.4.12             gdtools_0.5.2           
##  [37] enrichplot_1.32.0        ggrepel_0.9.8            tidytree_0.4.8          
##  [40] codetools_0.2-20         DelayedArray_0.38.2      ggforce_0.5.0           
##  [43] DOSE_4.6.0               tidyselect_1.2.1         ggseqlogo_0.2.2         
##  [46] aplot_0.3.2              locfdr_1.1-8             UCSC.utils_1.8.0        
##  [49] farver_2.1.2             BiocFileCache_3.2.0      GenomicAlignments_1.48.0
##  [52] jsonlite_2.0.0           ellipsis_0.3.3           systemfonts_1.3.2       
##  [55] tools_4.6.1              ggnewscale_0.5.2         progress_1.2.3          
##  [58] ragg_1.5.2               treeio_1.36.1            Rcpp_1.1.2              
##  [61] glue_1.8.1               SparseArray_1.12.3       xfun_0.62               
##  [64] qvalue_2.44.0            usethis_3.2.2            withr_3.0.3             
##  [67] BiocManager_1.30.27      fastmap_1.2.0            boot_1.3-32             
##  [70] callr_3.8.0              digest_0.6.39            R6_2.6.1                
##  [73] gridGraphics_0.5-1       textshaping_1.0.5        GO.db_3.23.1            
##  [76] biomaRt_2.68.0           RSQLite_3.53.3           cigarillo_1.2.1         
##  [79] utf8_1.2.6               tidyr_1.3.2              fontLiberation_0.1.0    
##  [82] htmlwidgets_1.6.4        prettyunits_1.2.0        httr_1.4.9              
##  [85] S4Arrays_1.12.1          scatterpie_0.2.6         pkgconfig_2.0.3         
##  [88] gtable_0.3.6             blob_1.3.0               S7_0.2.2                
##  [91] clusterProfiler_4.20.3   brio_1.1.5               htmltools_0.5.9         
##  [94] fontBitstreamVera_0.1.1  ProtGenerics_1.44.0      scales_1.4.0            
##  [97] png_0.1-9                ggfun_0.2.2              knitr_1.52              
## [100] rstudioapi_0.19.0        reshape2_1.4.5           rjson_0.2.23            
## [103] nlme_3.1-169             curl_8.1.0               org.Hs.eg.db_3.23.1     
## [106] cachem_1.1.0             stringr_1.6.0            BiocVersion_3.23.1      
## [109] parallel_4.6.1           restfulr_0.0.17          desc_1.4.3              
## [112] pillar_1.11.1            grid_4.6.1               vctrs_0.7.3             
## [115] tidydr_0.0.7             dbplyr_2.6.0             cluster_2.1.8.2         
## [118] tximport_1.40.0          evaluate_1.0.5           cli_3.6.6               
## [121] locfit_1.5-9.12          compiler_4.6.1           Rsamtools_2.28.0        
## [124] rlang_1.3.0              crayon_1.5.3             labeling_0.4.3          
## [127] plyr_1.8.9               fs_2.1.0                 ggiraph_0.9.6           
## [130] stringi_1.8.9            BiocParallel_1.46.0      txdbmaker_1.8.0         
## [133] lazyeval_0.2.3           devtools_2.5.2           fontquiver_0.2.1        
## [136] GOSemSim_2.38.3          Matrix_1.7-5             hms_1.1.4               
## [139] patchwork_1.3.2          bit64_4.8.6              KEGGREST_1.52.2         
## [142] statmod_1.5.2            AnnotationHub_4.2.2      igraph_2.3.4            
## [145] memoise_2.0.1            bslib_0.12.0             ggtree_4.2.0            
## [148] bit_4.6.0                gson_0.2.2               ape_5.8-1
```
