## Supplementary material for "Deducing splicing events from transcript sets with splicelogic": Novel transcripts workflow: NovelTxps_workflow.html


### Novel Transcripts Workflow

```
library(splicelogic)
library(SummarizedExperiment)
library(GenomicFeatures)
library(dplyr)
library(ggplot2)
library(tibble)
library(here)
library(plyxp)
# splicelogic devel, loaded from source: clone the repo next to this one
#   git clone -b devel https://github.com/thelovelab/splicelogic.git ../splicelogic
# then load it with devtools (install.packages("devtools") if needed)
devtools::load_all(here("../splicelogic"))
```

#### Load bambu output

`bambu_discovery.rds` is the exon-by-transcript
`GRangesList` from bambu’s discovery run (novel plus
annotated transcripts); `bambu_quant_se.rds` is the matching
quantification `SummarizedExperiment`.

```
bdisc    <- readRDS(here("workflows", "data", "bambu_discovery.rds"))
se_bambu <- readRDS(here("workflows", "data", "bambu_quant_se.rds"))
bdisc
```

```
## GRangesList object of length 390071:
## $BambuTx1
## GRanges object with 2 ranges and 2 metadata columns:
##       seqnames        ranges strand | exon_rank exon_endRank
##          <Rle>     <IRanges>  <Rle> | <integer>    <integer>
##   [1]     chr1 840205-840361      + |         1            2
##   [2]     chr1 840434-840538      + |         2            1
##   -------
##   seqinfo: 3366 sequences from an unspecified genome; no seqlengths
## 
## $BambuTx2
## GRanges object with 2 ranges and 2 metadata columns:
##       seqnames          ranges strand | exon_rank exon_endRank
##          <Rle>       <IRanges>  <Rle> | <integer>    <integer>
##   [1]     chr1 1352691-1353159      + |         1            2
##   [2]     chr1 1353270-1353573      + |         2            1
##   -------
##   seqinfo: 3366 sequences from an unspecified genome; no seqlengths
## 
## $BambuTx3
## GRanges object with 3 ranges and 2 metadata columns:
##       seqnames          ranges strand | exon_rank exon_endRank
##          <Rle>       <IRanges>  <Rle> | <integer>    <integer>
##   [1]     chr1 1615449-1615633      + |         1            3
##   [2]     chr1 1616508-1616614      + |         2            2
##   [3]     chr1 1617075-1617323      + |         3            1
##   -------
##   seqinfo: 3366 sequences from an unspecified genome; no seqlengths
## 
## ...
## <390068 more elements>
```

#### Transcript annotation (condition, CPM, direction)

Condition and donor live only in the sample names; `ctrl`
-> WT and `exp` -> KD to match the satuRn workflow. For
each transcript we take the mean CPM in each group and the
`log2(WT / KD)` fold change, whose sign gives the
direction.

```
cn <- colnames(se_bambu)
colData(se_bambu)$condition <- factor(
  ifelse(grepl("_ctrl", cn), "WT", "KD"), levels = c("KD", "WT")
)
colData(se_bambu)$donor <- sub("-0008.*$", "", cn)
table(colData(se_bambu)$donor, colData(se_bambu)$condition)
```

```
##            
##             KD WT
##   GTEX-QEG5  1  1
##   GTEX-QV44  1  2
##   GTEX-S4Z8  2  1
##   GTEX-S95S  2  1
```

```
cpm   <- assay(se_bambu, "CPM")
is_wt <- colData(se_bambu)$condition == "WT"

rowData(se_bambu)$cpm_WT <- rowMeans(cpm[,  is_wt, drop = FALSE])
rowData(se_bambu)$cpm_KD <- rowMeans(cpm[, !is_wt, drop = FALSE])
rowData(se_bambu)$lfc_WT_vs_KD <- log2((rowData(se_bambu)$cpm_WT + 1) /
                                       (rowData(se_bambu)$cpm_KD + 1))
rowData(se_bambu)$direction <- ifelse(
  rowData(se_bambu)$cpm_WT > rowData(se_bambu)$cpm_KD, "up_in_WT",
  ifelse(rowData(se_bambu)$cpm_WT < rowData(se_bambu)$cpm_KD, "up_in_KD", "tie")
)

# most `tie` rows are transcripts at zero in both groups
table(rowData(se_bambu)$direction)
```

```
## 
##      tie up_in_KD up_in_WT 
##   236725    72419    80927
```

```
table(rowData(se_bambu)$direction[rowMeans(cpm) > 1])
```

```
## 
## up_in_KD up_in_WT 
##    12511    14875
```

How many samples each donor contributes to each condition.

```
samples_per_donor <- as.data.frame(colData(se_bambu)) |>
  dplyr::count(donor, condition) |>
  tidyr::pivot_wider(names_from = condition, values_from = n, values_fill = 0) |>
  dplyr::mutate(total = WT + KD)

# with a total row
dplyr::bind_rows(
  samples_per_donor,
  dplyr::summarise(samples_per_donor, donor = "total",
                   dplyr::across(c(KD, WT, total), sum))
)
```

```
## # A tibble: 5 × 4
##   donor        KD    WT total
##   <chr>     <int> <int> <int>
## 1 GTEX-QEG5     1     1     2
## 2 GTEX-QV44     1     2     3
## 3 GTEX-S4Z8     2     1     3
## 4 GTEX-S95S     2     1     3
## 5 total         6     5    11
```

What the bambu objects record about the run. Neither stores the bambu
version, and bambu builds its `SummarizedExperiment` itself,
so tximeta is not involved. The discovery object keeps the novel
discovery rate (NDR) threshold bambu used.

```
names(metadata(se_bambu))
```

```
## [1] "incompatibleCounts" "warnings"
```

```
names(metadata(bdisc))
```

```
## [1] "NDRthreshold" "warnings"
```

```
metadata(bdisc)$NDRthreshold
```

```
## [1] 0.244
```

A quick look at the novel genes’ expression, to see how much signal
there is.

```
xp <- se_bambu |> new_plyxp()
xp_novel <- xp |>
  filter(rows(novelGene)) |>
  mutate(rows(mean_cpm = rowMeans(.assays_asis$CPM)))
nrow(xp_novel)
```

```
## [1] 2815
```

```
table(cut(log10(rowData(xp_novel)$mean_cpm), -3:3))
```

```
## 
## (-3,-2] (-2,-1]  (-1,0]   (0,1]   (1,2]   (2,3] 
##       1     390    1430     804     167      22
```

```
hist(log10(rowData(xp_novel)$mean_cpm))
```

##### Expression filter

Keep transcripts with raw count >= 10 in at least 6 samples.
Applied before building the exon object so both the novel
(`up`) and annotated (`down`) partitions are
filtered.

```
# transcripts per type, before and after the filter
count_tx_type <- \(se) table(
  tx_type = ifelse(rowData(se)$novelTranscript, "novel", "annotated")
)
n_before <- count_tx_type(se_bambu)

min_count   <- 10
min_samples <- 6
keep_counts <- rowSums(assay(se_bambu, "counts") >= min_count) >= min_samples
se_bambu    <- se_bambu[keep_counts, ]
dim(se_bambu)
```

```
## [1] 20916    11
```

```
n_after <- count_tx_type(se_bambu)
```

```
filter_counts <- dplyr::bind_rows(
  "unfiltered" = as.data.frame(n_before, responseName = "n"),
  "filtered"   = as.data.frame(n_after,  responseName = "n"),
  .id = "stage"
) |>
  dplyr::mutate(stage = factor(stage, levels = c("unfiltered", "filtered"))) |>
  dplyr::group_by(tx_type) |>
  # the filtered bar also shows the share retained
  dplyr::mutate(label = ifelse(
    stage == "filtered",
    sprintf("%s\n(%.1f%%)", scales::comma(n), 100 * n / max(n)),
    scales::comma(n)
  ))

ggplot(filter_counts, aes(x = tx_type, y = n, fill = stage)) +
  geom_col(position = position_dodge(width = 0.8), width = 0.7) +
  geom_text(aes(label = label), position = position_dodge(width = 0.8),
            vjust = -0.3, size = 3.2, lineheight = 0.9, colour = "grey20") +
  scale_fill_manual(values = c(
    "unfiltered" = "#B0CFFF",  # light blue
    "filtered"   = "#3A86FF"   # darker blue
  )) +
  # log scale from 1: the annotated set is ~100x the novel set, so on a
  # linear axis the novel bars would be slivers. Every decade is labelled.
  scale_y_log10(breaks = 10^(0:6), minor_breaks = NULL,
                labels = scales::comma, limits = c(1, NA),
                expand = expansion(mult = c(0, 0.12))) +
  labs(title = "Transcripts before and after the expression filter",
       subtitle = sprintf("Filter: raw count ≥ %d in at least %d of %d samples",
                          min_count, min_samples, ncol(se_bambu)),
       x = NULL, y = "log(Number of Transcripts)", fill = NULL) +
  theme_bw() +
  theme(legend.position = "top")
```

##### Flat exon GRanges

A flat `GRanges`, one range per exon, with the
transcript-level annotation joined on. This is the exon object the
splicelogic `find_*()` and `plot_*()` functions
take.

```
exons <- unlist(bdisc, use.names = TRUE)
exons$tx_id <- names(exons)

# bdisc carries every discovered transcript; keep only those retained by the
# expression filter on se_bambu, so the match below is complete.
rd    <- rowData(se_bambu)
exons <- exons[exons$tx_id %in% rd$TXNAME]
i     <- match(exons$tx_id, rd$TXNAME)
stopifnot(!anyNA(i))

cols    <- c("GENEID", "novelGene", "novelTranscript", "txClassDescription",
             "NDR", "readCount", "relReadCount",
             "cpm_WT", "cpm_KD", "lfc_WT_vs_KD", "direction")
tx_info <- rd[, cols]
tx_info$mean_cpm <- rowMeans(assay(se_bambu, "CPM"))
tx_info$n_det    <- rowSums(assay(se_bambu, "counts") > 0)

mcols(exons) <- cbind(mcols(exons), tx_info[i, ])
names(mcols(exons))[names(mcols(exons)) == "GENEID"] <- "gene_id"

# bambu exons carry no exon_id. Build one from gene + coordinates (per the
# splicelogic vignette), so the same exon gets the same id in every transcript
# that contains it.
exons$exon_id <- paste0(exons$gene_id, ":",
                        start(exons), "-", end(exons))
```

##### Novel transcript counts

The comparison keeps novel transcripts at **annotated**
loci (so there is an annotated partner to compare against) and drops
novel transcripts at entirely new loci.

```
count_gr <- function(gr, label) {
  data.frame(set   = label,
             exons = length(gr),
             tx    = length(unique(gr$tx_id)),
             genes = length(unique(gr$gene_id)))
}

rbind(
  count_gr(exons, "all (extended annotation)"),
  count_gr(exons[exons$novelTranscript],  "novel transcripts"),
  count_gr(exons[!exons$novelTranscript], "annotated transcripts"),
  count_gr(exons[exons$novelTranscript &  exons$novelGene], "  novel tx, NEW locus (drop)"),
  count_gr(exons[exons$novelTranscript & !exons$novelGene], "  novel tx, annotated gene (keep)")
) |> print(row.names = FALSE)
```

```
##                                set  exons    tx genes
##          all (extended annotation) 216821 20916 12009
##                  novel transcripts   6295  1069   879
##              annotated transcripts 210526 19847 11387
##         novel tx, NEW locus (drop)   4118   699   541
##    novel tx, annotated gene (keep)   2177   370   338
```

#### Splice event detection with splicelogic

The two partitions are defined explicitly (no DTU test): the
**up** set is the novel transcripts at annotated loci, the
**down** (reference) set is the annotated transcripts.
`prepare_exons_by_partition()` assigns
`estimate = +1` to up and `-1` to down.

```
up   <- exons |> dplyr::filter(!novelGene, novelTranscript)
down <- exons |> dplyr::filter(!novelGene, !novelTranscript)

# genes present in both partitions are the ones with a comparison to make
length(intersect(unique(up$gene_id), unique(down$gene_id)))
```

```
## [1] 257
```

```
sl_exons <- prepare_exons_by_partition(up, down) |>
  preprocess(coef_col = "estimate")

# save sl_exons 
saveRDS(sl_exons, here("workflows", "data", "sl_exons_bambu.rds"))

events_bambu <- find_all_events(sl_exons)
```

```
## Calculating skipped exon events...
```

```
## Calculating included exon events...
```

```
## Calculating mutually exclusive exon events...
```

```
## Calculating retained intron events...
```

```
## Calculating spliced intron events...
```

```
## Calculating alternative 5' splice site events...
```

```
## Calculating alternative 3' splice site events...
```

```
## Calculating alternative transcription start site events...
```

```
## Calculating alternative transcription end site events...
```

```
## Done! 1181 events detected.
```

##### Collapse redundant events

An exon paired with several partner transcripts is reported once per
partner, as repeated rows with the same range. Collapsing those per
event type shows how much of the raw event count is that redundancy.
Rows are collapsed only for the same exon (gene, chromosome, start, end,
strand) and the same event type, as in the other workflows.

```
# define a helper function
my_paste <- \(x) paste(unique(x), collapse = ",")

# the exon is identified by `exon_key` (gene, chromosome, start, end and
# strand), built as in the vignette's recipe for an exon id, rather than by
# `exon_id`: the annotation can give the same exon more than one id
collapsed_events <- events_bambu |>
  dplyr::mutate(exon_key = paste(gene_id, seqnames, start, end, strand, sep = ":")) |>
  dplyr::group_by(exon_key, event_type) |>
  plyranges::reduce_ranges_directed(
    n           = plyranges::n(),
    exon_id     = my_paste(exon_id),
    tx_id       = my_paste(tx_id),
    exon_rank   = my_paste(exon_rank),
    event_tx_id = my_paste(event_tx_id)
  )
```

```
event_counts <- events_bambu |>
  as_tibble() |>
  dplyr::count(event_type, name = "count")

collapsed_event_counts <- collapsed_events |>
  as_tibble() |>
  dplyr::count(event_type, name = "count")

# combine both count tables into one, labelled by source
counts <- dplyr::bind_rows(
  "All events" = event_counts,
  "Collapsed events" = collapsed_event_counts,
  .id = "set"
)

# event types from most to fewest events
counts$event_type <- factor(
  counts$event_type,
  levels = event_counts$event_type[order(-event_counts$count)]
)

ggplot(counts, aes(x = event_type, y = count, fill = set)) +
  geom_col(position = position_dodge(width = 0.8)) +
  scale_fill_manual(values = c(
    "All events" = "#B0CFFF",       # light blue
    "Collapsed events" = "#3A86FF"  # darker blue
  )) +
  labs(title = "Novel-transcript splice event counts",
       x = "Event Type", y = "Count", fill = NULL) +
  theme_bw()
```

```
# Collapsed events by type, split into two groups: the transcript ends (ATSS,
# ATES) and the internal splicing events (everything else). One panel per
# group, bars in the package's event colours so each type matches its colour
# in plot_transcripts(), sorted by count within the group.
type_colors <- event_palette()
end_types   <- c("atss", "ates")

bar_counts <- collapsed_event_counts |>
  dplyr::mutate(
    pct   = 100 * count / sum(count),
    group = ifelse(event_type %in% end_types, "Transcript ends", "Internal splicing"),
    label = sprintf("%d (%.1f%%)", count, pct)
  )

# one panel per group, its height proportional to the number of types in it
plot_group <- function(grp, x_max, show_x) {
  d <- bar_counts |>
    dplyr::filter(group == grp) |>
    dplyr::arrange(count) |>
    dplyr::mutate(type = factor(toupper(event_type), levels = toupper(event_type)))
  ggplot(d, aes(count, type, fill = event_type)) +
    geom_col(width = 0.7) +
    geom_text(aes(label = label), hjust = -0.1, colour = "grey20", size = 3) +
    scale_fill_manual(values = type_colors, guide = "none") +
    scale_x_continuous(limits = c(0, x_max), expand = expansion(mult = c(0, 0.02))) +
    labs(title = sprintf("%s: %d (%.1f%%)", grp, sum(d$count), sum(d$pct)),
         x = if (show_x) "Events" else NULL, y = NULL) +
    theme_bw() +
    theme(plot.title = element_text(face = "bold", size = 10),
          panel.grid.major.y = element_blank(),
          panel.grid.minor = element_blank(),
          axis.text.x = if (show_x) element_text() else element_blank(),
          axis.ticks.x = if (show_x) element_line() else element_blank())
}

n_types <- table(bar_counts$group)

# the two panels stacked, with the overall title; the shared x axis runs past
# the longest bar to leave room for its count label
x_max <- max(bar_counts$count) * 1.35
patchwork::wrap_plots(
  plot_group("Transcript ends", x_max, show_x = FALSE),
  plot_group("Internal splicing", x_max, show_x = TRUE),
  ncol = 1,
  heights = c(n_types[["Transcript ends"]], n_types[["Internal splicing"]])
) +
  patchwork::plot_annotation(
    title = "Collapsed novel-transcript events by type",
    subtitle = sprintf("n = %s", scales::comma(sum(bar_counts$count)))
  )
```

A summary of the novel transcripts before and after the expression
filter, the genes they fall in, and the events they give.

```
# transcript annotation before the filter (se_bambu above is the filtered one)
rd_unfilt <- as.data.frame(rowData(readRDS(here("workflows", "data", "bambu_quant_se.rds"))))
rd_filt   <- as.data.frame(rowData(se_bambu))

tx_summary <- function(rd) {
  novel <- rd[rd$novelTranscript, ]
  c(novel_tx          = nrow(novel),
    novel_tx_known    = sum(!novel$novelGene),
    novel_tx_new_gene = sum(novel$novelGene),
    known_genes       = dplyr::n_distinct(novel$GENEID[!novel$novelGene]),
    new_genes         = dplyr::n_distinct(novel$GENEID[novel$novelGene]))
}
before <- tx_summary(rd_unfilt)
after  <- tx_summary(rd_filt)

# genes with a novel transcript and an annotated partner that passed the filter
compared_genes <- length(intersect(unique(up$gene_id), unique(down$gene_id)))

tibble::tibble(
  what = c("novel transcripts",
           "  at annotated genes (kept)",
           "  at new genes (dropped)",
           "annotated genes with a novel transcript",
           "  with an annotated partner after the filter",
           "new genes",
           "events (one row per transcript pair)",
           "distinct events (collapsed)"),
  before_filter = c(before[["novel_tx"]], before[["novel_tx_known"]],
                    before[["novel_tx_new_gene"]], before[["known_genes"]],
                    NA, before[["new_genes"]], NA, NA),
  after_filter  = c(after[["novel_tx"]], after[["novel_tx_known"]],
                    after[["novel_tx_new_gene"]], after[["known_genes"]],
                    compared_genes, after[["new_genes"]],
                    length(events_bambu), length(collapsed_events))
)
```

```
## # A tibble: 8 × 3
##   what                                           before_filter after_filter
##   <chr>                                                  <int>        <int>
## 1 "novel transcripts"                                     4402         1069
## 2 "  at annotated genes (kept)"                           1587          370
## 3 "  at new genes (dropped)"                              2815          699
## 4 "annotated genes with a novel transcript"               1372          338
## 5 "  with an annotated partner after the filter"            NA          257
## 6 "new genes"                                             1977          541
## 7 "events (one row per transcript pair)"                    NA         1181
## 8 "distinct events (collapsed)"                             NA          790
```

#### Per-transcript annotation for the plots

One label per transcript (novel status, then the direction and the
two group means) and a matching `log2(WT / KD)` value used to
draw the LFC strip beside each transcript.

```
# Pulled into a plain data.frame and read with with(): repeating `rd$` inside
# one call makes the console's `$` completion eat the paste.
d <- as.data.frame(rd[, c("TXNAME", "novelTranscript",
                          "cpm_WT", "cpm_KD", "lfc_WT_vs_KD")])

# novel/annot status, plus the direction and group means. A direction on two
# near-zero means is rounding noise, not a change, so those read "n.d.".
tx_annot <- with(d, paste0(
  ifelse(novelTranscript, "novel", "annot"), " | ",
  ifelse(cpm_WT + cpm_KD < 0.1, "n.d.",
         sprintf("log2 %+.2f  (WT %.2f / KD %.2f)", lfc_WT_vs_KD, cpm_WT, cpm_KD))
))
names(tx_annot) <- d[["TXNAME"]]

# log2(WT/KD) for the LFC strip: NA (no bar) where both groups are near zero.
tx_lfc <- with(d, ifelse(cpm_WT + cpm_KD < 0.1, NA_real_, lfc_WT_vs_KD))
names(tx_lfc) <- d[["TXNAME"]]

# one shared scale so bar lengths mean the same thing on every page
lfc_max <- quantile(abs(tx_lfc), 0.99, na.rm = TRUE)
```

#### Plotting helpers

Two small helpers that the package does not provide: a multi-page PDF
writer, and a per-gene layout that places
`plot_transcripts()` next to the companion
`plot_transcript_lfc()` strip (the package draws them as
separate plots; here `patchwork` composes them so they share
row order and y limits).

```
fig_dir <- here("workflows", "figures")
dir.create(fig_dir, showWarnings = FALSE)

# write a list of plots to one multi-page PDF, one plot per page
write_pdf <- function(plots, file, width = 10, height = 6) {
  grDevices::pdf(file, width = width, height = height, onefile = TRUE)
  on.exit(grDevices::dev.off(), add = TRUE)
  for (nm in names(plots)) print(plots[[nm]])
  invisible(file)
}

# plot_transcripts() + plot_transcript_lfc() side by side, one pair per gene
plot_gene_with_lfc <- function(events, exons, tx_lfc, lfc_max = NULL,
                               tx_annot = NULL, lfc_ends = c("KD", "WT"),
                               widths = c(5, 1.2), ...) {
  genes <- unique(as.character(events$gene_id))
  plots <- lapply(stats::setNames(genes, genes), function(g) {
    ev <- events[as.character(events$gene_id) == g]
    tryCatch({
      p <- plot_transcripts(ev, exons, title = g, tx_annot = tx_annot, ...)
      q <- plot_transcript_lfc(ev, tx_lfc, lfc_max = lfc_max,
                               lfc_ends = lfc_ends)
      patchwork::wrap_plots(p, q, widths = widths)
    }, error = function(e) {
      warning("Skipping ", g, ": ", conditionMessage(e))
      NULL
    })
  })
  Filter(Negate(is.null), plots)
}
```

#### Events for one gene

`exons` is already the flat `GRanges` the
package plotting functions take (one range per exon with a
`tx_id` column), so it is passed directly.

```
gene_of_interest <- unique(as.character(events_bambu$gene_id))[10]
events_for_gene  <- events_bambu[events_bambu$gene_id == gene_of_interest]

p <- plot_transcripts(events_for_gene, exons, title = gene_of_interest,
                      tx_annot = tx_annot)
q <- plot_transcript_lfc(events_for_gene, tx_lfc, lfc_max = lfc_max,
                         lfc_ends = c("KD", "WT"))
patchwork::wrap_plots(p, q, widths = c(5, 1.2))
```

#### All events, one page per gene (with the LFC strip)

```
gene_plots <- plot_gene_with_lfc(
  events_bambu, exons,
  tx_annot = tx_annot, tx_lfc = tx_lfc, lfc_max = lfc_max
)
length(gene_plots)

write_pdf(gene_plots, file.path(fig_dir, "events_by_gene_bambu.pdf"))
```

#### All events, one page per transcript pair

`plot_transcripts_by_pair()` splits on the transcript pair
internally, so `events_bambu` works directly.

```
pair_plots <- plot_transcripts_by_pair(events_bambu, exons)
length(pair_plots)

write_pdf(pair_plots, file.path(fig_dir, "events_by_pair_bambu.pdf"), height = 3)
```

#### Novel transcripts split by direction

Plot the novel transcripts separately by the direction of their WT/KD
change. In each event row the novel transcript is the one from the
`up` partition (`estimate > 0`):
`tx_id` when the row’s own estimate is positive, otherwise
its partner `event_tx_id`.

```
ev <- as.data.frame(S4Vectors::mcols(events_bambu)[, c("tx_id", "event_tx_id",
                                                       "estimate", "gene_id")])
novel_tx  <- with(ev, ifelse(estimate > 0, as.character(tx_id),
                             as.character(event_tx_id)))
novel_lfc <- unname(tx_lfc[novel_tx])   # NA when the novel transcript is n.d.

# which() drops the NA rows, so n.d. novel transcripts are in neither set
events_novel_kd <- events_bambu[which(novel_lfc < 0)]
events_novel_wt <- events_bambu[which(novel_lfc > 0)]

n_genes <- function(keep) length(unique(ev[["gene_id"]][keep]))
data.frame(
  set    = c("all events", "novel tx higher in KD (< 0)",
             "novel tx higher in WT (> 0)", "novel tx n.d. (not plotted)"),
  events = c(nrow(ev), sum(novel_lfc < 0, na.rm = TRUE),
             sum(novel_lfc > 0, na.rm = TRUE), sum(is.na(novel_lfc))),
  genes  = c(n_genes(TRUE), n_genes(which(novel_lfc < 0)),
             n_genes(which(novel_lfc > 0)), n_genes(is.na(novel_lfc)))
)
```

```
##                           set events genes
## 1                  all events   1181   257
## 2 novel tx higher in KD (< 0)    631   138
## 3 novel tx higher in WT (> 0)    550   127
## 4 novel tx n.d. (not plotted)      0     0
```

```
novel_sets <- list(novel_higher_in_KD = events_novel_kd,
                   novel_higher_in_WT = events_novel_wt)
```

A gene with novel transcripts in both directions appears in both
PDFs, each page showing only the novel transcripts of that direction and
their partners.

```
for (set in names(novel_sets)) {
  set_plots <- plot_gene_with_lfc(
    novel_sets[[set]], exons,
    tx_annot = tx_annot, tx_lfc = tx_lfc, lfc_max = lfc_max
  )
  set_file <- file.path(fig_dir, paste0("events_", set, "_bambu.pdf"))
  write_pdf(set_plots, set_file)
  message(set, ": ", length(set_plots), " pages written to ", set_file)
}
```

#### Splice-site logos for the novel-transcript events

Every event exon is a real spliced exon, so its flanking splice sites
should be the canonical ones: the **3’ acceptor**
(`...AG |` exon) in the region **upstream** of
the exon, and the **5’ donor** (exon `| GU...`)
**downstream**. This is a sanity check that the
novel-transcript events sit at genuine splice sites. A first exon has no
acceptor (the bases before it are the transcription start) and a last
exon no donor (the transcription end), so each side only uses the exons
that actually have that splice site. Many novel events sit on terminal
exons: most A5SS exons are first exons and most A3SS exons last ones.
ATSS and ATES events are included on the same rule: an ATSS exon is a
first exon, so only its donor is used, and an ATES exon a last exon, so
only its acceptor.

```
source(here("workflows/helper.R"))
```

```
# Restrict to the standard chromosomes: the bambu object carries hundreds of
# _fix/_alt scaffolds that are not all in the BSgenome, which otherwise triggers
# a noisy seqinfo-merge warning (and errors for exons on absent scaffolds).
# pruning.mode = "coarse" drops ranges on non-standard seqlevels.
standard_chr <- function(gr) {
  gr <- GenomeInfoDb::keepStandardChromosomes(gr, pruning.mode = "coarse")
  GenomeInfoDb::seqlevels(gr) <- GenomeInfoDb::seqlevelsInUse(gr)
  gr
}

# every event type, ATSS and ATES included: the first/last-exon rule below
# keeps only the donor of an ATSS exon and the acceptor of an ATES exon
splice_events <- standard_chr(events_bambu)

# each event row describes the transcript that contains the exon: novel (the
# up partition) or annotated (down). The annotated event exons are the SE
# exons and one side of each MXE pair; everything else is novel.
is_novel <- splice_events$novelTranscript

# One 1-bp range per distinct splice site: the first base of an exon marks its
# acceptor, the last base its donor (strand-aware). Flanking these gives the
# same intronic bases as flanking the whole exon, and a boundary shared by
# several transcripts counts once. Acceptors come from exons that are not
# first, donors from exons that are not last (bambu's exon_endRank is 1 on the
# last exon).
acceptor_sites <- function(gr) {
  gr <- GenomicRanges::granges(gr[gr$exon_rank > 1])
  unique(GenomicRanges::resize(gr, 1, fix = "start"))
}
donor_sites <- function(gr) {
  gr <- GenomicRanges::granges(gr[gr$exon_endRank > 1])
  unique(GenomicRanges::resize(gr, 1, fix = "end"))
}

# 10 bp into the intron on each side (exon_bases = 0, so purely intronic):
# the upstream logo should end in AG, the downstream logo should start with GU.
splice_site_mats <- function(gr) {
  acc <- slFlankLogoMatrices(acceptor_sites(gr), genome = "hg38",
                             width = 10, exon_bases = 0)
  don <- slFlankLogoMatrices(donor_sites(gr), genome = "hg38",
                             width = 10, exon_bases = 0)
  structure(list(upstream = acc$upstream, downstream = don$downstream),
            width = 10, exon_bases = 0,
            n_sites = c(upstream   = attr(acc, "n_exons"),
                        downstream = attr(don, "n_exons")))
}

plot_splice_site_logo <- function(mats, title) {
  n <- attr(mats, "n_sites")
  slPlotFlankLogo(mats, method = "bits", titles = c(
    upstream   = sprintf("Upstream: 3' splice site (acceptor), n = %d",
                         n[["upstream"]]),
    downstream = sprintf("Downstream: 5' splice site (donor), n = %d",
                         n[["downstream"]])
  )) +
    patchwork::plot_annotation(
      title   = title,
      caption = paste("distinct sites; acceptors from non-first exons,",
                      "donors from non-last exons")
    )
}

novel_logo <- splice_site_mats(splice_events[is_novel])
plot_splice_site_logo(novel_logo, "Event exons in novel transcripts")
```

As a reference, every exon of the annotated transcripts (the
`down` partition), not only the annotated event exons, which
are just the SE exons and one side of each MXE pair. These are GENCODE
exons, so they should be almost entirely canonical.

```
annotated_exons <- standard_chr(down)
annotated_logo  <- splice_site_mats(annotated_exons)
plot_splice_site_logo(annotated_logo, "All exons of annotated transcripts")
```

##### Non-canonical splice sites

To find the events that break the pattern, take the 2 intronic bases
immediately flanking each exon — strand-aware, via
`plyranges::flank_*()` — and read them with
`get_seq()`. The acceptor (upstream) should be
`AG`, the donor (downstream) `GU`. Anything else
is flagged. As for the logos, a first exon has no acceptor and a last
exon no donor; those are recorded as `NA` and not
checked.

```
# one row per distinct exon within each set, so an exon paired with several
# partners is not counted repeatedly
ex_once <- c(slUniqueExons(splice_events[is_novel]),
             slUniqueExons(splice_events[!is_novel]))
# an exon of a single-exon transcript has no splice site to check
ex_once <- ex_once[ex_once$exon_rank > 1 | ex_once$exon_endRank > 1]

# 2 bp immediately 5' (acceptor) and 3' (donor) of each exon, in transcription
# order; get_seq() reverse-complements minus-strand ranges automatically
acceptor <- ex_once |> plyranges::flank_upstream(width = 2)
donor    <- ex_once |> plyranges::flank_downstream(width = 2)

# NA where the exon has no such site: no acceptor before a first exon, no
# donor after a last one
ex_once$acceptor <- ifelse(
  ex_once$exon_rank > 1,
  as.character(get_seq(acceptor, "hg38", as_rna = TRUE)), NA
)
ex_once$donor <- ifelse(
  ex_once$exon_endRank > 1,
  as.character(get_seq(donor, "hg38", as_rna = TRUE)), NA
)
ex_once$canonical <- (is.na(ex_once$acceptor) | ex_once$acceptor == "AG") &
                     (is.na(ex_once$donor)    | ex_once$donor    == "GU")

# how many follow the canonical AG/GU pattern
table(canonical = ex_once$canonical)
```

```
## canonical
## FALSE  TRUE 
##    16   690
```

```
table(acceptor = ex_once$acceptor)
```

```
## acceptor
##  AC  AG 
##   4 430
```

```
table(donor = ex_once$donor)
```

```
## donor
##  AU  GC  GU 
##   6   6 441
```

```
table(novel = ex_once$novelTranscript, canonical = ex_once$canonical)
```

```
##        canonical
## novel   FALSE TRUE
##   FALSE     2  115
##   TRUE     14  575
```

The comparison below counts splice sites rather than exons, in two
sets: the splice sites of the novel event exons
(`splice_events`, every event type), against every splice
site of the annotated transcripts (`down`) as the reference.
Each distinct site counts once per set, with the same first/last-exon
rule as above, so an ATSS exon contributes only its donor and an ATES
exon only its acceptor.

```
# acceptor (AG) and donor (GU) dinucleotide of every distinct splice site
site_canonical <- function(gr) {
  read2 <- function(sites, flank) {
    if (length(sites) == 0L) return(character(0))
    as.character(get_seq(flank(sites, width = 2), "hg38", as_rna = TRUE))
  }
  acc <- read2(acceptor_sites(gr), plyranges::flank_upstream)
  don <- read2(donor_sites(gr),    plyranges::flank_downstream)
  dplyr::bind_rows(
    tibble::tibble(site = "acceptor", seq = acc, canonical = acc == "AG"),
    tibble::tibble(site = "donor",    seq = don, canonical = don == "GU")
  )
}

novel_events <- splice_events[is_novel]

site_sets <- dplyr::bind_rows(
  "annotated transcripts" = site_canonical(annotated_exons),
  "novel event exons"     = site_canonical(novel_events),
  .id = "set"
)

site_sets |>
  dplyr::count(set, site, canonical) |>
  tidyr::pivot_wider(names_from = canonical, values_from = n,
                     values_fill = 0L, names_prefix = "canonical_")
```

```
## # A tibble: 4 × 4
##   set                   site     canonical_FALSE canonical_TRUE
##   <chr>                 <chr>              <int>          <int>
## 1 annotated transcripts acceptor             183         114416
## 2 annotated transcripts donor               1121         114089
## 3 novel event exons     acceptor               2            313
## 4 novel event exons     donor                 12            346
```

```
# the novel event sites per event type: ATSS adds only donors, ATES only
# acceptors (a site shared by two event types is listed under both)
split(novel_events, novel_events$event_type) |>
  lapply(site_canonical) |>
  dplyr::bind_rows(.id = "event_type") |>
  dplyr::count(event_type, site, canonical) |>
  tidyr::pivot_wider(names_from = canonical, values_from = n,
                     values_fill = 0L, names_prefix = "canonical_")
```

```
## # A tibble: 12 × 4
##    event_type site     canonical_FALSE canonical_TRUE
##    <chr>      <chr>              <int>          <int>
##  1 a3ss       acceptor               1             42
##  2 a3ss       donor                  0             26
##  3 a5ss       acceptor               0             22
##  4 a5ss       donor                  6             59
##  5 ates       acceptor               2            210
##  6 atss       donor                 10            243
##  7 ie         acceptor               0             51
##  8 ie         donor                  0             51
##  9 mxe        acceptor               0              7
## 10 mxe        donor                  0              7
## 11 ri         acceptor               0              5
## 12 ri         donor                  0              3
```

```
canon_p <- fisher.test(table(site_sets$set, site_sets$canonical))$p.value
canon_p
```

```
## [1] 4.580079e-05
```

```
canon_counts <- site_sets |>
  dplyr::mutate(
    set  = factor(set, levels = c("annotated transcripts",
                                  "novel event exons")),
    site = factor(ifelse(canonical, "canonical (AG/GU)", "non-canonical"),
                  levels = c("non-canonical", "canonical (AG/GU)"))
  ) |>
  dplyr::count(set, site) |>
  # keep zero counts, so a set with no non-canonical sites still gets a label
  tidyr::complete(set, site, fill = list(n = 0L)) |>
  dplyr::group_by(set) |>
  dplyr::mutate(frac  = n / sum(n),
                label = sprintf("%s (%.1f%%)", scales::comma(n), 100 * frac),
                set   = sprintf("%s\n(n = %s)", set, scales::comma(sum(n)))) |>
  dplyr::ungroup() |>
  # annotated at the bottom, novel event exons on top
  dplyr::mutate(set = factor(set, levels = unique(set)))

ggplot(canon_counts, aes(x = frac, y = set, fill = site)) +
  # non-canonical share on the left
  geom_col(position = position_stack(reverse = TRUE), width = 0.6) +
  # non-canonical count just right of its segment, canonical at the bar's end
  geom_text(data = \(d) dplyr::filter(d, site == "non-canonical"),
            aes(label = label), hjust = 0, nudge_x = 0.01,
            size = 3.5, colour = "grey20") +
  geom_text(data = \(d) dplyr::filter(d, site == "canonical (AG/GU)"),
            aes(x = 0.99, label = label), hjust = 1,
            size = 3.5, colour = "grey20") +
  scale_fill_manual(values = c("non-canonical"     = "#3A86FF",
                               "canonical (AG/GU)" = "grey85")) +
  scale_x_continuous(labels = scales::percent, expand = c(0, 0)) +
  labs(title = "Canonical splice sites, novel event exons vs annotated",
       subtitle = sprintf(paste0("distinct splice sites, ATSS donors and ATES ",
                                 "acceptors included\nFisher's exact test p = %.2g"),
                          canon_p),
       x = "Share of splice sites", y = NULL, fill = NULL) +
  theme_bw() +
  theme(legend.position = "top",
        # room for the "100%" axis label at the right edge
        plot.margin = margin(5.5, 14, 5.5, 5.5))
```

The novel event exons on their own, counted as exons: an exon is
canonical when every splice site it has is (so an ATSS exon is judged on
its donor, an ATES exon on its acceptor).

```
novel_exon_counts <- ex_once[ex_once$novelTranscript] |>
  as_tibble() |>
  dplyr::mutate(site = factor(
    ifelse(canonical, "canonical (AG/GU)", "non-canonical"),
    levels = c("canonical (AG/GU)", "non-canonical")
  )) |>
  dplyr::count(site, .drop = FALSE) |>
  dplyr::mutate(label = sprintf("%d (%.1f%%)", n, 100 * n / sum(n)))

ggplot(novel_exon_counts, aes(x = site, y = n, fill = site)) +
  geom_col(width = 0.6) +
  geom_text(aes(label = label), vjust = -0.4, size = 3.5, colour = "grey20") +
  scale_fill_manual(values = c("non-canonical"     = "#3A86FF",
                               "canonical (AG/GU)" = "grey70"),
                    guide = "none") +
  scale_y_continuous(expand = expansion(mult = c(0, 0.1))) +
  labs(title = "Novel event exons",
       subtitle = sprintf("n = %d exons\nATSS donors and ATES acceptors included",
                          sum(novel_exon_counts$n)),
       x = NULL, y = "Exons") +
  theme_bw()
```

```
# the offenders: event exons whose acceptor is not AG or donor is not GU
noncanonical <- ex_once[!ex_once$canonical]

noncanonical |>
  as_tibble() |>
  dplyr::select(seqnames, start, end, strand, gene_id, tx_id,
                novelTranscript, exon_rank, event_type, acceptor, donor) |>
  dplyr::arrange(acceptor, donor) |>
  print(n = 20, width = Inf)
```

```
## # A tibble: 16 × 11
##    seqnames     start       end strand gene_id            tx_id            
##    <fct>        <int>     <int> <fct>  <chr>              <chr>            
##  1 chr13    113196181 113196222 -      ENSG00000126226.22 ENST00000337344.9
##  2 chr17     81682034  81683594 -      ENSG00000214087.10 ENST00000577142.1
##  3 chrM         14717     15652 +      ENSG00000198727.2  BambuTx4279      
##  4 chrM          4208      4267 +      ENSG00000210082.2  BambuTx914       
##  5 chr3     184243552 184243630 -      ENSG00000214160.10 BambuTx208       
##  6 chr17     81683691  81683763 -      ENSG00000214087.10 BambuTx756       
##  7 chr20      4023717   4024199 +      ENSG00000226608.3  BambuTx832       
##  8 chrM          1680      3996 +      ENSG00000210082.2  BambuTx914       
##  9 chr1      31908194  31908430 -      ENSG00000184007.22 BambuTx27        
## 10 chrM         13971     14482 +      ENSG00000198727.2  BambuTx4279      
## 11 chr5        443176    443254 +      ENSG00000180104.16 BambuTx2702      
## 12 chr5      93682978  93683570 -      ENSG00000249353.2  BambuTx3990      
## 13 chr6      35259525  35259727 +      ENSG00000065029.15 BambuTx299       
## 14 chrM         12797     13051 +      ENSG00000198786.2  BambuTx916       
## 15 chr5     163437595 163437652 +      ENSG00000113328.21 BambuTx2015      
## 16 chr9      19049416  19049536 +      ENSG00000155876.6  BambuTx2093      
##    novelTranscript exon_rank event_type acceptor donor
##    <lgl>               <int> <chr>      <chr>    <chr>
##  1 FALSE                   5 se         AC       GU   
##  2 FALSE                   2 si         AC       GU   
##  3 TRUE                    2 a3ss       AC       <NA> 
##  4 TRUE                    2 ates       AC       <NA> 
##  5 TRUE                    7 a5ss       AG       AU   
##  6 TRUE                    1 a5ss       <NA>     AU   
##  7 TRUE                    1 a5ss       <NA>     AU   
##  8 TRUE                    1 a5ss       <NA>     AU   
##  9 TRUE                    1 atss       <NA>     AU   
## 10 TRUE                    1 atss       <NA>     AU   
## 11 TRUE                    1 a5ss       <NA>     GC   
## 12 TRUE                    1 a5ss       <NA>     GC   
## 13 TRUE                    1 atss       <NA>     GC   
## 14 TRUE                    1 atss       <NA>     GC   
## 15 TRUE                    1 atss       <NA>     GC   
## 16 TRUE                    1 atss       <NA>     GC
```

To eyeball one, read the whole exon with 5 intronic bases on each
side, in transcription order (`get_seq()` reverse-complements
the minus strand). The acceptor is the 2 intronic bases just before the
exon (canonical `AG`), the donor the 2 intronic bases just
after it (canonical `GU`):

```
one <- noncanonical[1]

up5   <- get_seq(plyranges::flank_upstream(one,   width = 5), "hg38", as_rna = TRUE)
exon  <- get_seq(one,                                         "hg38", as_rna = TRUE)
down5 <- get_seq(plyranges::flank_downstream(one, width = 5), "hg38", as_rna = TRUE)

# intron(5) | EXON | intron(5), 5'->3'
cat(sprintf("%s | %s | %s\n", up5, exon, down5))
```

```
## UCCAG | GACAGGGGAAAGUAUCUUGUCGCUGCUGAGGGAUCCCUCCAAAAGGAAGGUUCCACCCCAGCCCCUUACACCCAACCAU | AUCCU
```

```
data.frame(
  intron_up_5   = as.character(up5),   # ...ends in AG for a canonical acceptor
  exon          = as.character(exon),
  intron_down_5 = as.character(down5)  # starts with GU for a canonical donor
)
```

```
##   intron_up_5
## 1       UCCAG
##                                                                              exon
## 1 GACAGGGGAAAGUAUCUUGUCGCUGCUGAGGGAUCCCUCCAAAAGGAAGGUUCCACCCCAGCCCCUUACACCCAACCAU
##   intron_down_5
## 1         AUCCU
```

#### Session info

The R and package versions (satuRn, splicelogic, Bioconductor, …)
this report was knitted with.

```
sessionInfo()
```

```
## R version 4.6.1 (2026-06-24)
## Platform: aarch64-apple-darwin23
## Running under: macOS Sequoia 15.6.1
## 
## Matrix products: default
## BLAS:   /Library/Frameworks/R.framework/Versions/4.6/Resources/lib/libRblas.0.dylib 
## LAPACK: /Library/Frameworks/R.framework/Versions/4.6/Resources/lib/libRlapack.dylib;  LAPACK version 3.12.1
## 
## locale:
## [1] en_US.UTF-8/en_US.UTF-8/en_US.UTF-8/C/en_US.UTF-8/en_US.UTF-8
## 
## time zone: America/New_York
## tzcode source: internal
## 
## attached base packages:
## [1] stats4    stats     graphics  grDevices utils     datasets  methods  
## [8] base     
## 
## other attached packages:
##  [1] BSgenome.Hsapiens.UCSC.hg38_1.4.5 BSgenome_1.80.0                  
##  [3] rtracklayer_1.72.0                BiocIO_1.22.0                    
##  [5] Biostrings_2.80.2                 XVector_0.52.0                   
##  [7] GenomeInfoDb_1.48.0               splicelogic_1.1.4                
##  [9] testthat_3.3.2                    plyranges_1.32.0                 
## [11] plyxp_1.6.1                       here_1.0.2                       
## [13] tibble_3.3.1                      ggplot2_4.0.3                    
## [15] dplyr_1.2.1                       GenomicFeatures_1.64.0           
## [17] AnnotationDbi_1.74.0              SummarizedExperiment_1.42.0      
## [19] Biobase_2.72.0                    GenomicRanges_1.64.0             
## [21] Seqinfo_1.2.0                     IRanges_2.46.0                   
## [23] S4Vectors_0.50.3                  BiocGenerics_0.58.1              
## [25] generics_0.1.4                    MatrixGenerics_1.24.0            
## [27] matrixStats_1.5.0                
## 
## loaded via a namespace (and not attached):
##  [1] tidyselect_1.2.1         farver_2.1.2             blob_1.3.0              
##  [4] S7_0.2.2                 bitops_1.1-0             fastmap_1.2.0           
##  [7] RCurl_1.98-1.20          GenomicAlignments_1.48.0 XML_3.99-0.25           
## [10] digest_0.6.39            lifecycle_1.0.5          ellipsis_0.3.3          
## [13] KEGGREST_1.52.2          RSQLite_3.53.3           magrittr_2.0.5          
## [16] compiler_4.6.1           rlang_1.3.0              sass_0.4.10             
## [19] tools_4.6.1              utf8_1.2.6               yaml_2.3.12             
## [22] knitr_1.52               labeling_0.4.3           S4Arrays_1.12.1         
## [25] bit_4.6.0                pkgbuild_1.4.8           curl_8.1.0              
## [28] DelayedArray_0.38.2      RColorBrewer_1.1-3       pkgload_1.5.3           
## [31] abind_1.4-8              BiocParallel_1.46.0      withr_3.0.3             
## [34] purrr_1.2.2              desc_1.4.3               grid_4.6.1              
## [37] scales_1.4.0             cli_3.6.6                rmarkdown_2.32          
## [40] crayon_1.5.3             otel_0.2.0               rstudioapi_0.19.0       
## [43] httr_1.4.9               rjson_0.2.23             sessioninfo_1.2.4       
## [46] DBI_1.3.0                cachem_1.1.0             ggseqlogo_0.2.2         
## [49] parallel_4.6.1           restfulr_0.0.17          vctrs_0.7.3             
## [52] devtools_2.5.2           Matrix_1.7-5             jsonlite_2.0.0          
## [55] patchwork_1.3.2          bit64_4.8.6              jquerylib_0.1.4         
## [58] tidyr_1.3.2              glue_1.8.1               codetools_0.2-20        
## [61] gtable_0.3.6             UCSC.utils_1.8.0         pillar_1.11.1           
## [64] brio_1.1.5               htmltools_0.5.9          R6_2.6.1                
## [67] rprojroot_2.1.1          evaluate_1.0.5           lattice_0.22-9          
## [70] png_0.1-9                Rsamtools_2.28.0         cigarillo_1.2.1         
## [73] memoise_2.0.1            bslib_0.12.0             SparseArray_1.12.3      
## [76] xfun_0.62                usethis_3.2.2            fs_2.1.0                
## [79] pkgconfig_2.0.3
```
